# METTL5, an 18S rRNA-specific m^6^A methyltransferase, modulates expression of stress response genes

**DOI:** 10.1101/2020.04.27.064162

**Authors:** Hao Chen, Qi Liu, Dan Yu, Kundhavai Natchiar, Chen Zhou, Chih-hung Hsu, Pang-Hung Hsu, Xing Zhang, Bruno Klaholz, Richard I. Gregory, Xiaodong Cheng, Yang Shi

## Abstract

RNA N^6^-methyladenosine (m^6^A) modification is present in different RNA molecules, including protein-coding mRNAs and non-coding RNAs such as ribosomal RNAs (rRNAs). Previous studies identified m^6^A in both the 18S and 28S rRNAs, but the roles of these methylation events are poorly understood due to the lack of knowledge of the responsible methyltransferases. Here, we report that mammalian METTL5, a member of a highly conserved methyltransferase family, specifically methylates adenosine 1832 (A1832) in the 18S rRNA *in vivo* and *in vitro*. We identify TRMT112 as a near stoichiometric partner of METTL5 important for the enzymatic activity of METTL5. By mapping the positions of translating ribosomes (Ribo-seq), we found translation of multiple stress response-related mRNAs, including Atf4 mRNA, is selectively reduced in the Mettl5 knockout (KO) mouse B16 melanoma cells. Atf4 is a key transcription factor that mediates the Integrated Stress Response (ISR), as exemplified by the Endoplasmic Reticulum (ER) stress. Consistently, transcription of ISR effector genes is reduced in Mettl5 KO cells during ER stress, suggesting a compromised ISR. Our findings reveal a new mechanism that regulates expression of stress response genes and suggest that chemical modifications of ribosomal RNAs may play a key role in selectively impacting translation and possibly ISR.

## Introduction

RNAs are heavily modified by chemical modifications. To date, more than 100 chemically modified nucleotides have been documented in different types of RNA in all kingdoms of life (1). Among them, N^6^-methyladenosine (m^6^A) is present in many RNA types from bacteria to mammal and plays vital roles in determining the messenger RNA (mRNA) fates. Functional studies suggest that m^6^A in mRNAs impacts cellular processes of RNA biology, including pre-mRNA splicing, nuclear transport, mRNA stability and translational initiation (2–4). Besides mRNA, m^6^A is also present in the functionally important regions in the ribosomal RNA (rRNA) of both the small and large subunits of the ribosome (5,6).

The ribosome is one of the most conserved and sophisticated molecular machines in the cell. For human rRNAs, there are more than 200 residues that are enzymatically modified in the fully mature ribosome (5,7). The modified residues are mainly located in regions of the rRNAs that are known to be functionally important (5,7,8). Therefore, identification of enzymes responsible for rRNA modifications is expected to not only shed mechanistic light on translational regulation but also provide guidance for the design of selective inhibitors to probe mechanisms and to potentially treat human diseases.

Recently, ZCCHC4 was identified as the enzyme responsible for m^6^A methylation of A4220 in the 28S rRNA and shown to be important for ribosomal maturation and global translation (9). Analysis of the high-resolution cryogenic electron microscopy structures of ribosome demonstrate that, in addition to 28S rRNA, a site in the 18S rRNA, A1832, is also methylated at the N^6^ position (5,6). A1832 is proximal to the mRNA channel in the decoding center, which underlines the potential importance of methylation of this site in translational regulation and calls for an identification and characterization of its methyltransferase.

Protein synthesis is indispensable for the basic function of cell and its regulation plays a crucial role in the overall response of cell to stress (10,11). In eukaryotic cells, mRNAs mainly employ a cap-dependent scanning mechanism to initiate translation (12,13). After the mRNA translation initiation factor, eIF4E, recognizes the m^7^G cap structure, a 43S pre-initiation complex (PIC) is formed and scans the mRNA to search for a proper start codon (AUG) to initiate translation. If an upstream open reading frame (uORF) is short, a portion of the small ribosomal subunit remains attached to mRNA following termination of uORF translation, then resumes scanning the same mRNA and re-initiates translation at a downstream site. The translational re-initiation is highly regulated and represents an important mechanism for controlling translation of stress response genes, such as the ATF4 transcription factor (14–16). Indeed, ATF4 is at the nexus of multiple signaling pathways that regulate responses of cells to various types of stress, emanating from “sensor” kinases to phosphorylation of translational regulators such as eIF2α, which leads to activation of ATF4 expression at the translational level (17). This kinase-eIF2α-ATF4 axis is proposed to be a major mechanism that mediates the Integrated Stress Response (ISR), which is typified by the endoplasmic reticulum (ER) stress (18,19).

ATF4 translation is regulated through the alternative usage of its two different uORFs: uORF1 and uORF2 (17,20,21). While uORF1 positively facilitates ribosome scanning and re-initiation of translation at a downstream coding region of the ATF4 mRNA, leading to productive translation of ATF4 (17,20), uORF2 is in a different reading frame and functions as an inhibitory element to suppress ATF4 translation (17). In non-stressed cells, uORF2 is preferentially translated, which blocks ATF4 expression (17,20). Under stress conditions, however, “sensor” kinases phosphorylate eIF2α, leading to bypass of uORF2 and preferential use of ATF4 downstream coding sequence (CDS). Consequently, this results in a productive translation of ATF4, which induces transcription of many more stress responsive factors (17,20). Previous studies of ATF4 translational control were focused on the signaling regulators such as the kinases for eIF2α. More recently, a study demonstrated that m^6^A modification in uORF2 of ATF4 mRNA could control ATF4 translation, probably through recruiting m^6^A reader proteins or changing the secondary structure of mRNA (21). However, whether the core machinery of translation, i.e., the ribosome itself, plays a role in ATF4 translational control and ISR remains largely unknown.

To address the above questions, we set out to identify the methyltransferase responsible for m^6^A methylation of A1832 in the 18S rRNA. We provide multiple lines of evidence that mammalian METTL5, a member of the conserved METTL family of methyltransferases (22), mediates 18S rRNA m^6^A methylation *in vitro* and *in vivo*. We employed Ribo-seq to identify genome-wide translational changes in response to Mettl5 knockout (KO) and found that stress-responsive mRNAs are more significantly affected by the loss of Mettl5 and 18S rRNA m^6^A methylation in mouse B16 melanoma cells. Importantly, translational re-initiation of Atf4 CDS is significantly diminished as a result of Mettl5 KO, suggesting a compromised ISR. Our investigations not only discover and define the enzymological properties of the 18S rRNA m^6^A metyltransferase, METTL5, but also suggest that 18S rRNA m^6^A methylation may represent a new mechanism that modulates translation of stress-related mRNAs, including that of ATF4, important for cellular stress response.

## Results

### METTL5 mediates m^6^A methylation of 18S rRNA

18S rRNA has been shown to be methylated at A1832 (5,6). To identify the methyltransferase that mediates A1832 methylation, we carried out a mini-screen of candidate enzymes, all of which contain a Rossmann-fold domain (23) with the [DNSH]PP[YFW] motif (24,25) shown to be important for the catalytic activity for some members such as METTL3 (26) (Fig. S1A). Compared with METTL3, METTL4, METTL16 and ZCCHC4, only METTL5 knockdown induces the most significant decrease of m^6^A level in 18S rRNA in human 293T cells. Mass spectrometry analysis shows that less than 30% of m^6^A of 18S rRNA remained after METTL5 knockdown (KD) (Fig. 1A and Fig. S1B). To further confirm that METTL5 is responsible for installing m^6^A in the 18S rRNA, we generated a METTL5 KO 293T cell line, in which exon3 of *METTL5* in both alleles was deleted and a translational frame-shift was created. Compared with METTL5 KD, we observed an even more striking reduction of 18S rRNA m^6^A in the METTL5 KO cells where almost all 18S rRNA methylation was abrogated (Fig. 1B). As a control, m^6^A in the 28S rRNA, which is mediated by ZCCHC4 (9), is unaffected in the METTL5 KO cells. Importantly, 18S m^6^A is restored by re-expression of wildtype but not by the catalytically inactive form of METTL5 (N_126_PPF_129_ motif mutated to APPA) (Fig. 1B and Fig. S1C). In agreement with its role in modifying ribosomal RNAs, we found METTL5 predominantly localizes to the nucleolus, which is the site of ribosome biogenesis (Fig. S1D). Using phylogenetic analysis, we identified METTL5 homologs in metazoa and plants, but not in yeast and prokaryotes (Fig. 1C). Importantly, the presence of a considerable amount of N^6^-adenosine methylation in the small subunit (SSU), but not the large subunit (LSU), of rRNA is correlated with the presence of METTL5 (Fig. 1D), consistent with METTL5 being the m^6^A methyltransferase for the small subunit of rRNA.

**Figure 1.**
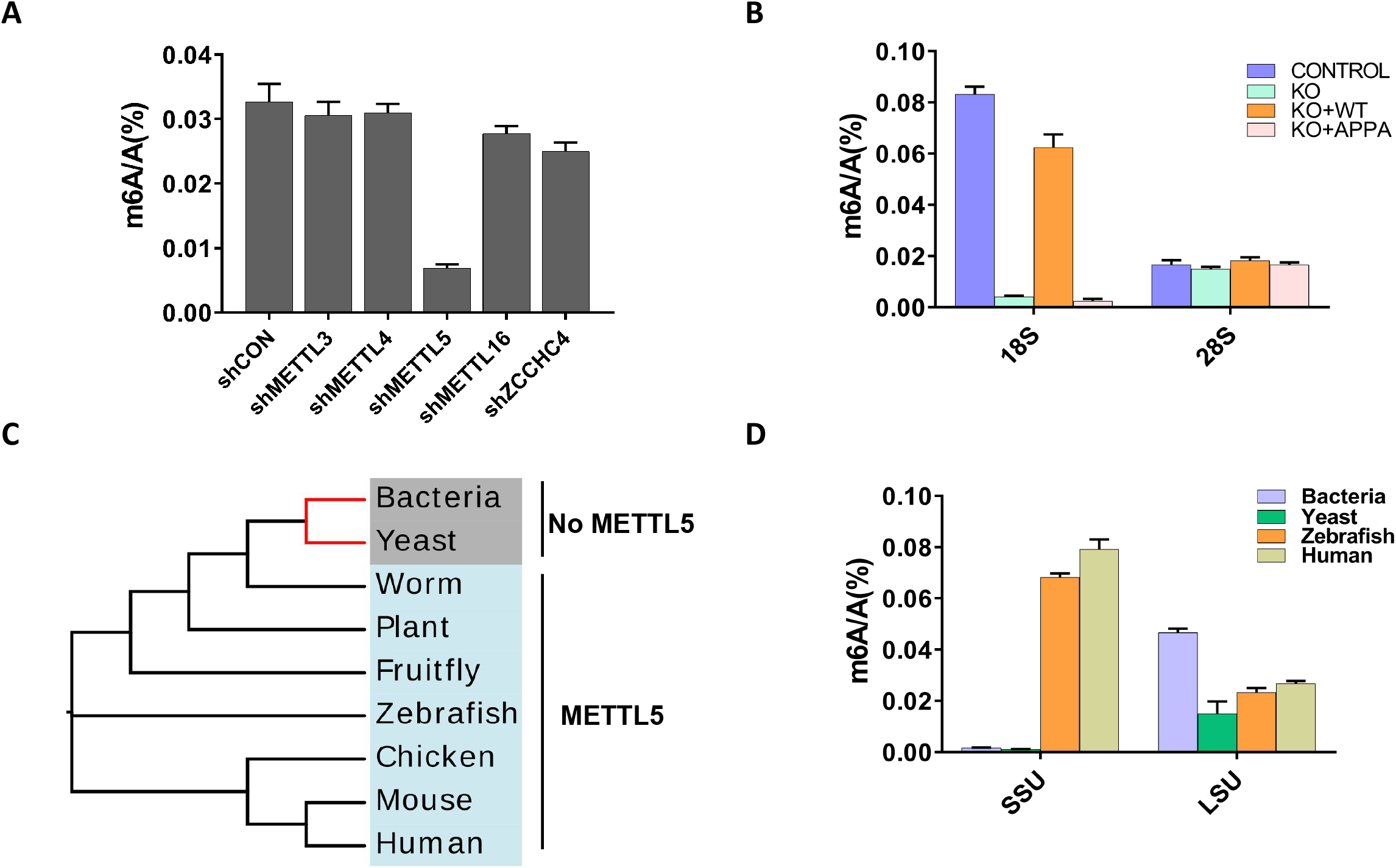
METTL5 is involved in m^6^A methylation of 18S rRNA. A. LC–MS/MS showing that depletion of METTL5 in 293T cells, but not other known and putative m^6^A methyltransferases, leads to a drastic decrease of m^6^A in 18S rRNA. n = 3 independent experiments; B. Re-expression of WT, but not a catalytic mutant METTL5^APPA^, in METTL5 KO 293T cell lines restored m^6^A modification of 18S rRNA but has no significant impact on 28S m^6^A level; C. Phylogenetic analysis demonstrating that METTL5 is an evolutionarily conserved m^6^A methyltransferase from worm to human, but absent in many lower species, including bacteria and yeast; D. LC–MS/MS quantification of m^6^A in rRNA demonstrating that considerable amount of small subunit of rRNA m^6^A can be detected only in species carrying the METTL5 gene.

### TRMT112 is a partner of METTL5 and is important for METTL5 activity

Within the METTL family of methyltransferases, METTL3 mediates catalysis of m^6^A on mRNA, but the active methyltransferase is a complex of METTL3 and METTL14 (26,27). To investigate whether METTL5 functions in a similar manner, we carried out tandem affinity purification using FLAG and HA-tagged METTL5 (METTL5-FLAG-HA). Commassie blue staining of the purified FLAG-HA-tagged METTL5 from 293T stable cells identified a single prominent band associated with METTL5 (Fig. 2A), which was shown to be TRMT12 by Western blotting (Fig. S2A), which is consistent with the previous report (31). Furthermore, untagged, endogenous METTL5 also co-immunoprecipitates (co-IP) TRMT112, and reciprocally, TRMT112 antibodies brought down considerable amount of endogenous METTL5 (Fig. 2B&C). TRMT112 has been identified as a co-factor for multiple methyltransferases, including those that modify rRNA, tRNA and proteins (28–30). As shown in Fig. 2D, although bacterially purified recombinant METTL5 alone is able to mediate 18S rRNA methylation, this ability is significantly enhanced (~100 fold higher) when TRMT112 was added into the reaction, suggesting that METTL5 and TRMT112 may function together to specifically catalyze 18S rRNA m^6^A methylation. Next, we determined the kinetics of recombinant METTL5-TRMT112 complex (co-expressed and co-purified from *E. coli*; Fig. S2C) on RNA oligo derived from 18S rRNA and a control GACU oligo (a known substrate of METTL3/14 complex). As expected, the METTL5-TRMT112 complex has much stronger activity towards the 18S rRNA oligo compared with the control oligo (*k*_cat_ = 17.1 h^−1^ vs <1 h^−1^) (Fig. 2E), consistent with the notion that METTL5-TRMT112 specifically mediates methylation of 18S rRNA. A recent independent study also reported TRMT112 as METTL5 partner, and that the METTL5 catalytic activity *in vivo* requires TRMT112 (31). Somewhat surprisingly, van Tran et al. failed to observe *in vitro* activity of the METTL5-TRMT112 heterodimer on short RNA oligonucleotides corresponding to the sequences surrounding A1832 in the mature ribosome (31).

**Figure 2.**
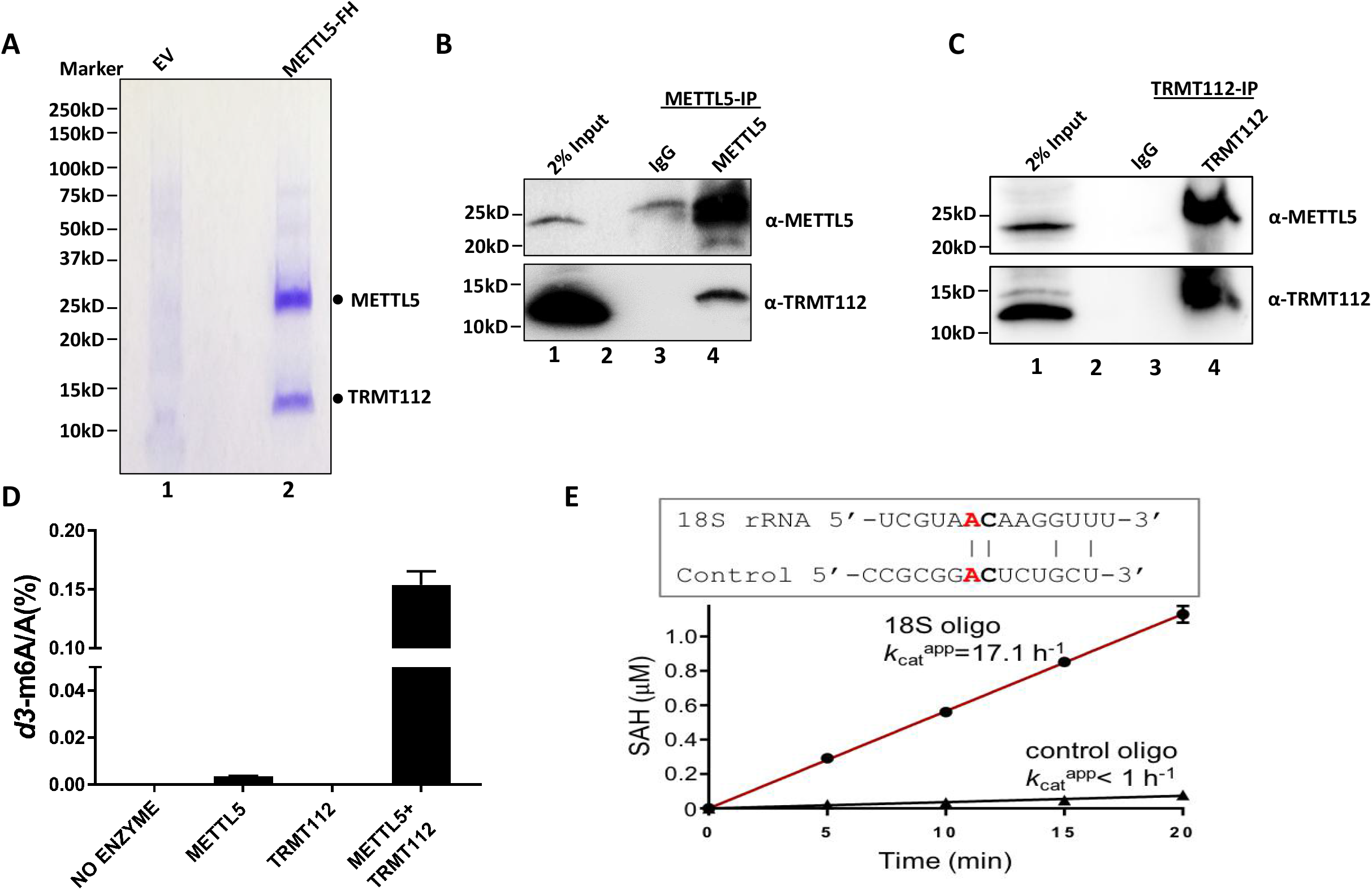
TRMT112 is a critical partner of METTL5 and important for its enzymatic activity. A. METTL5 complex was obtained by FLAG and HA tandem purification from FLAG-HA-tagged METTL5-expressing stable cell lines. The Coomassie Blue Staining shows that the complex contains an enriched band at around 12 kD, which was shown to be TRMT112; EV: empty vector; B&C. Reciprocal co-IP experiments indicating that endogenous METTL5 interacts with TRMT112; D. In vitro methylation assay of recombinant METTL5 protein on 18S rRNA. METTL5 enzymatic activity was significantly enhanced when TRMT112 was added into the reaction. n = 2 independent experiments; E. Recombinant METTL5-TRMT112 complex is active on RNA oligo derived from 18S. Comparison of a 14-nt RNA oligo from 18S rRNA and a control oligo, which shares AC dinucleotides.

### METTL5 makes contacts with nucleic acids neighboring A1832 in 18S rRNA

Next, we used enhanced CLIP (eCLIP) (32) to identify METTL5-bound RNAs, which may represent potential substrates for METTL5. METTL5-bound RNAs covalently linked by UV irradiation were extracted for RNA sequencing. We find that rRNA fragments were the most abundant RNAs bound to METTL5, but other types of RNAs identified by eCLIP are relatively low (Fig. S3). Notably, the strongest signals identified by METTL5 eCLIP are proximal to and appear to surround A1832 of 18S rRNA based on the three-dimensional structure of the human ribosome (Fig. 3A and 3B). To verify that METTL5 is specifically targeted to this region, we isolated the 18S rRNA fragment using biotin labeled 36-nt DNA oligo complementary to the A1832 region (detailed in Methods). HPLC-MS/MS analysis showed that the m^6^A signal is noticeably higher surrounding A1832 compared with full-length 18S rRNA (Fig. 3C, full-length (FL) 18S rRNA in WT *vs* enriched A1832-neighboring fragment in WT). As a control, the same experiment done in METTL5 KO cells found no enrichment of m^6^A signals (Fig. 3C). Thus, these findings support our hypothesis that METTL5 is the main methyltransferase responsible for A1832 methylation in human 18S rRNA and A1832 is a major site methylated by METTL5.

**Figure 3.**
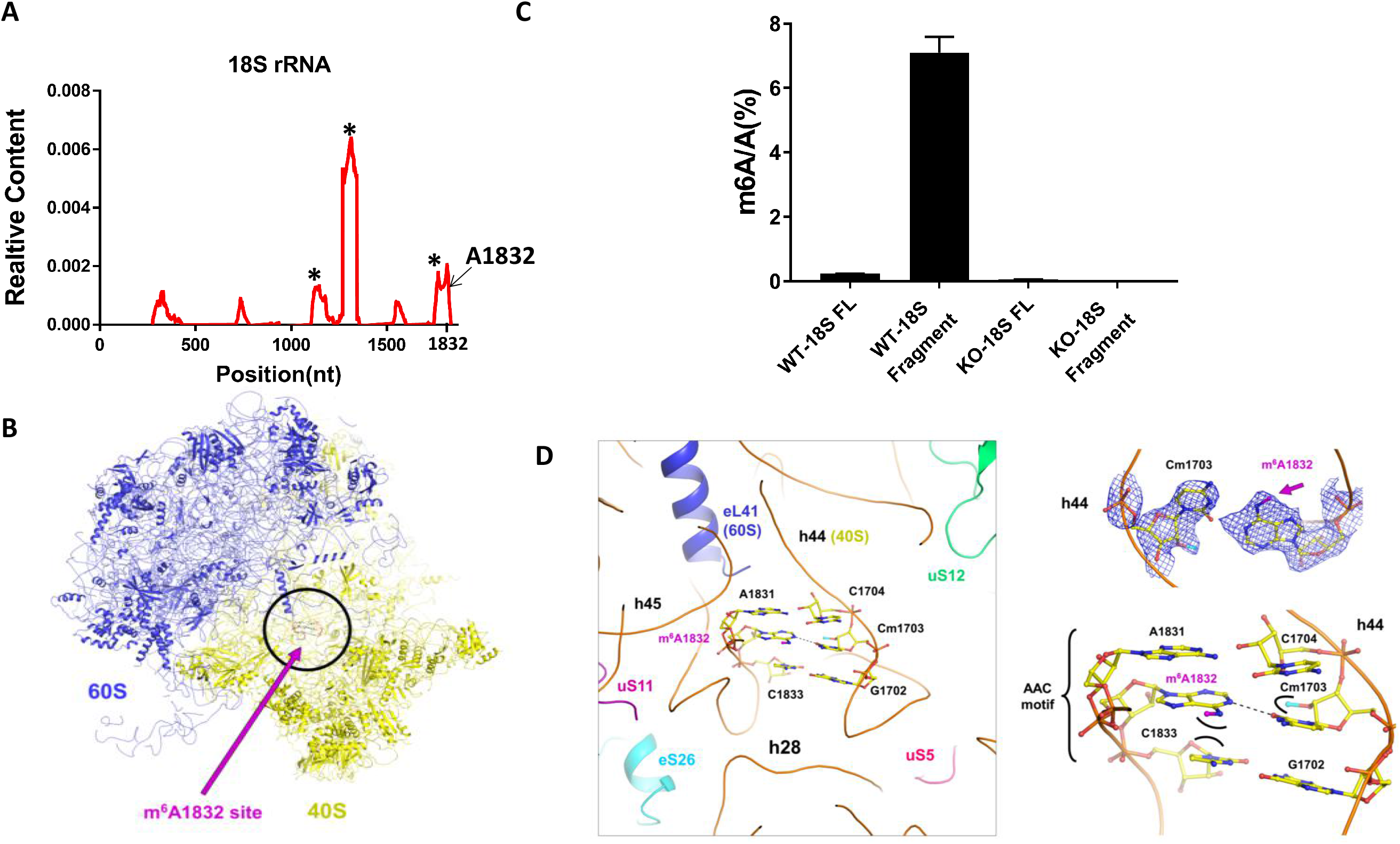
METTL5 specifically binds 18S rRNA regions surrounding A1832. A&B. eCLIP experiments identifying peaks of 18S rRNA representing contact sites of METTL5 are mainly localized in the A1832-centered region in the 3’ terminus of 18S rRNA. Notably, many of the strongest signals identified by METTL5 eCLIP are proximal to and appear to surround A1832 of 18S rRNA based on the three-dimensional structure of the human ribosome, and the peaks surround A1832 in 3D structure are indicated with asterisks in (A). n = 2 independent replicates; C. Cryo-EM structure shows that the interaction of m^6^A1832 with neighboring nucleotides is important for proper structure of ribosome; D. HPLC-MS/MS results indicating that the RNA fragment neighboring the A1832 is the major m^6^A region in 18S rRNA. n = 3 independent replicates.

### Mettl5 depletion decreases translation of stress response-related mRNAs

We next analyzed the high-resolution Cryo-EM structure of human 80□S ribosome and found that A1832 of 18S rRNA is close to the decoding center and N^6^-methyl group in A1832 is predicted to increase the base stacking with the neighboring bases (A1831 and C1833) (Fig. 3D). Interestingly, the partner base (C1703) is, in addition, also 2’-O methylated, which on the complementary strand helps stabilize base stacking between Cm1703 and its neighbors. The modified site is at a place with a structural transition from a classical C1833-G1702 pairing to a non-canonical A1832-C1703 (which is intrinsically less stable) and a non-interacting A1831-C1704 (see the AAC base triplet in Fig. 3C), in which the m^6^A1832 is involved in stacking with the neighboring C1833 and A1831. Furthermore, it is interesting to note that the site is on helix h44, a key region in the 40S subunit next to the decoding center, close to the A1824/A1825 nucleotides, which are essential for tRNA decoding. These structural observations imply that the m^6^A modification catalyzed by METTL5 may affect translation activity (Fig. 3D). To address this hypothesis, we investigated mRNA translation status in the presence and absence of METTL5 by carrying out sucrose density gradient to obtain ribosome profiles. Surprisingly, we found no detectable differences in terms of global distributions of mono- and poly-ribosomes between WT and METTL5 KO 293T cells (Fig. S4A). This finding suggests that 18S rRNA methylation is unlikely to influence ribosomal biogenesis or basal activity of translation, but might represent a fine-tuning mechanism to regulate translation of specific RNAs or translation under specific biological conditions.

To experimentally test this hypothesis, we employed Ribosome Profiling coupled with deep sequencing (Ribo-seq) (33,34) to obtain insight into translation at the genome-wide scale in mouse B16 melanoma cells. The initial quality control step showed that Ribo-seq data quality met the requirement for downstream analysis (Fig. S4B and S4C). Global mapping of translation sites and quantitative measurement of the ribosome density at individual mRNAs identified 277 translationally altered genes among 10,466 actively transcribed genes between WT and KO cells (Fig. 4A and Fig. S4D) with more down-regulated (183) than up-regulated genes (94). Bioinformatics analysis found that the down-regulated genes appear to have lower minimum free energy (MFE) throughout the entire mRNA (including 5’ UTR, coding region and 3’UTR) compared with non-regulated genes (Fig. 4B), while there are no significant differences in codon (except CCC and CGU) usage between WT and KO B16 cells(Fig. S4E). Gene Ontology (GO) analysis of the 183 down-regulated genes retrieved GO terms related to stress response, apoptosis and translational regulation (Fig. 4C). Similarly, Gene Set Enrichment Analysis (GSEA) also points to the involvement of Mettl5 in the regulation of stress response (Fig. 4D). In contrast, we didn’t find significantly enriched pathways for the up-regulated genes. Together, these results suggest that Mettl5-mediated 18S m^6^A methylation plays important roles in controlling translation of specific genes, but this methylation event does not appear to be essential for ribosomal maturation or global translation activity.

**Figure 4.**
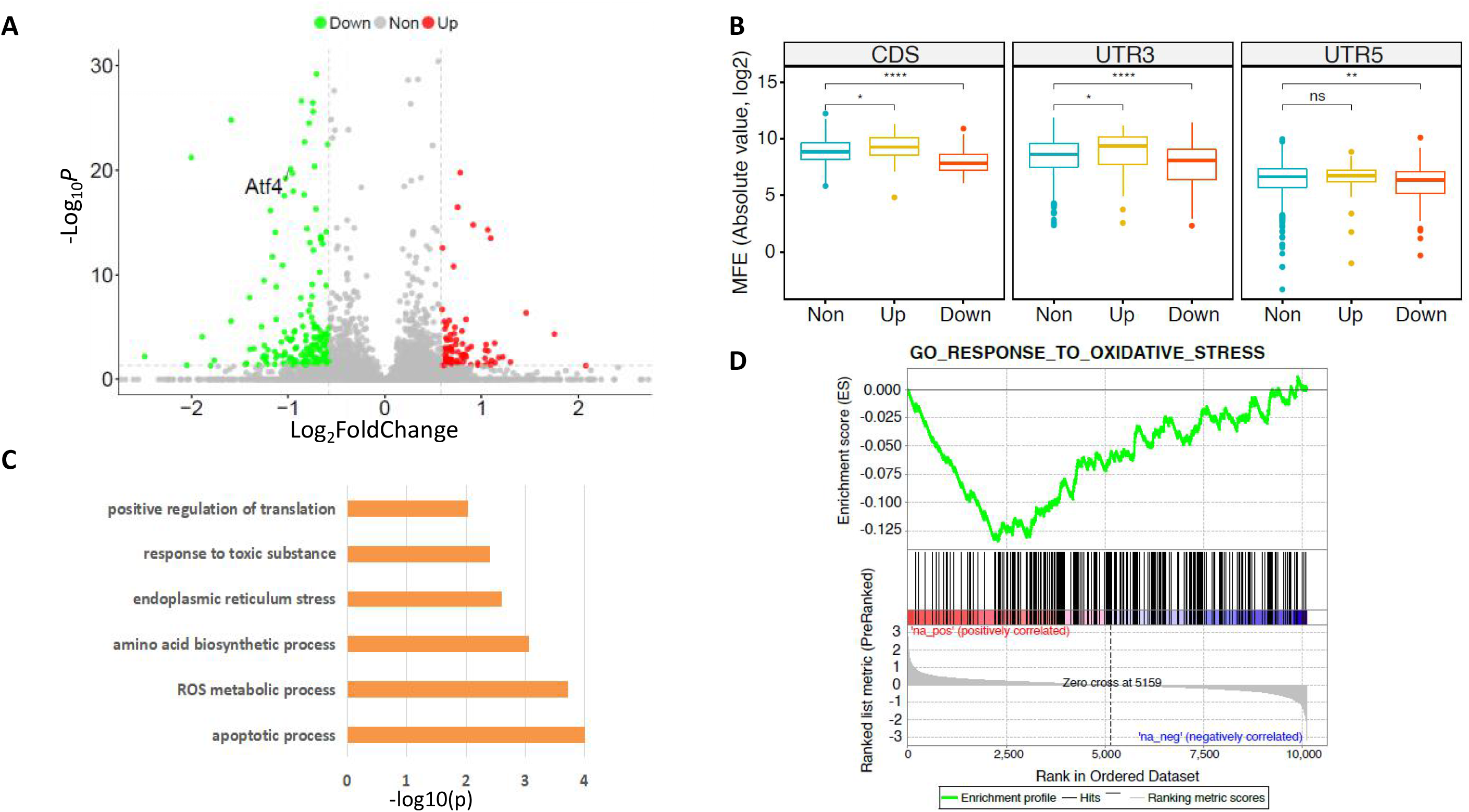
Mettl5 depletion diminishes translation of stress-responsive genes. A. Translation of 277 genes identified to be significantly affected by depletion of Mettl5 in B16 cells, cut-off > 1.5-fold changes, n=2 independent experiments; B. Ribo-seq revealing that translation of mRNAs with lower MFE levels is more significantly diminished by Mettl5 KO in B16 cells; C. GO analysis identifying the down-regulated genes being enriched in pathways related to stress response; D. GSEA analysis identifying the affected genes to be enriched in the stress-related pathways.

### Mettl5 is necessary for the productive translation of Atf4 through alternate uORF usage during the Integrated Stress Response (ISR)

Among the 277 significantly altered genes in response to Mettl5 KO, the transcription factor Atf4 is of particular interest due to its established role in the ISR (15). The ISR is primarily mediated by the eIF2α signaling pathways, culminating in bypassing of the repressive uORF2 and activation of productive Atf4 translation. Atf4 subsequently directs specific gene expression programs to mount an ISR, which is a critical mechanism employed by both normal and tumor cells to survive various types of stress (17). Ribo-seq showed that translation of uORF1 (Fig. 5A, highlighted in blue) and Atf4 CDS significantly decreases in the Mettl5 KO cells, but the negative cis-regulatory element uORF2 (Fig. 5A, highlighted in pink) remains by and large unaffected. To validate this observation, we separated the monosomes and polysomes (which are associated with actively translated mRNAs) by sucrose gradient and determined the distribution of Atf4 mRNAs in polysomes by RT-qPCR. Consistently, Atf4 mRNA is significantly less enriched in the polysome fractions upon Mettl5 KO (Fig. 5B), An Atf4 translation reporter system (Vattem and Wek, 2004) further confirmed that Atf4 translation is indeed reduced in the Mettl5 KO cells (Fig. 5C), suggesting that 18S rRNA methylation by Mettl5 regulates productive translation of Atf4.

**Figure 5.**
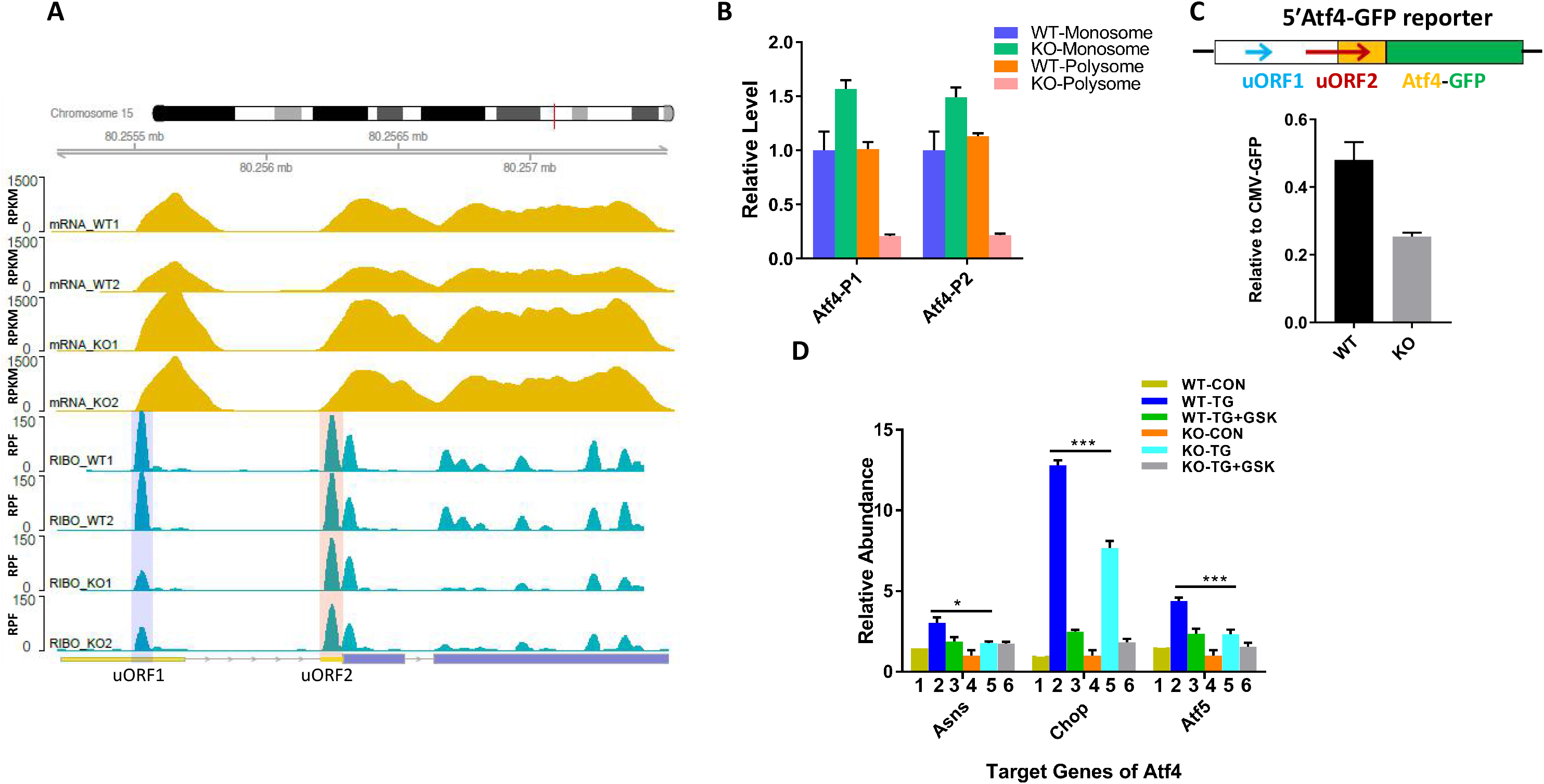
Mettl5 is necessary for efficient translation of Atf4 during the Integrated Stress Response (ISR). A. Translation of Atf4 uORF1 and mORF, but not uORF2, is disrupted upon Mettl5 KO in B16 cells. The uORF1 and uORF2 regions were highlighted with blue and pink color, respectively; B. Mettl5 depletion in B16 cells reduces the distribution of Atf4 mRNA in the polysome fractions (number of ribosomes: 3 and greater), which are associated with active translation. Atf4-P1 and Atf4-P2 represent two distinct primer sets for detecting Atf4 levels; C. Atf4-GFP reporter assay showing that Atf4 translation is inhibited upon Mettl5 KO in B16 cells. The density was normalized by CMV-driven expression of GFP; D. The transcription of Atf4 target genes (*Asns*, *Chop* and *Atf5*) is compromised in Mettl5 KO B16 cells due to a reduced Atf4 level, *: p-value < 0.05, ***: p-value < 0.001.

Given the role of Atf4 production in the ISR, we next determined whether loss of Mettl5 might impair the ISR. Importantly, we found that in response to ER stress induced by Thapsigargin (TG) (17), transcription of *Asns*, *Chop* and *Atf5*, which are well-characterized Atf4 target genes, is also reduced in Mettl5 KO cells in response to ER stress induced by TG (Fig. 5D, compare 2 and 5 in the TG treatment groups). Furthermore, the differences in mRNA levels of Atf4 target genes were still largely observed (at least for *Chop* and *Atf5*) after treatment by the Perk kinase (Protein kinase RNA-like endoplasmic reticulum kinase) inhibitor GSK2606414 (Fig. 5D, compare 3 and 6 in the TG treatment groups), suggesting that Mettl5 and eIF2α-phosphorylation may represent two parallel pathways that regulate cellular stress responses (Fig. 5D), although additional investigations are necessary to provide a complete understanding of this observation at both functional and mechanistic levels. Taken together, the above results support the hypothesis that Mettl5 may plays a role in regulating ISR via controlling Atf4 translation.

## Discussion

rRNA methylation is believed to play a role in ribosome biogenesis and maturation (7). Importantly, we identify a new, regulatory role for ribosomal RNA methylation in the expression of stress response genes. Our conclusion is supported by multiple lines of evidence. First, we demonstrate that METTL5 specifically mediates methylation of 18S rRNA at A1832 *in vitro* and *in vivo*. Second, we show that removal of A1832 methylation by knocking out Mettl5 only compromises translation of a subset of mRNAs in mouse B16 melanoma cells. GO analysis showed that mRNAs affected by Mettl5 belong to genes involved in stress responses such as ISR, in which the transcription factor Atf4 plays a major role. Mettl5 null cells are compromised in their ability to initiate productive Atf4 translation and are therefore more sensitive to ISR stress. Our findings thus highlight a previously unappreciated role of rRNA methylation in regulating mRNA translation rather than ribosome biogenesis and also reveal a novel mechanism of Atf4 regulation important for ISR.

### Transition from m^4^C to m^6^A modification in 18S rRNA during evolution

The Cryo-EM structure showed that 18S rRNA A1832 is partially base-paired with C1702 and localized in region of helix 44 (h44), which holds mRNA and decodes mRNA into protein (5). This region has been reported to be intensively decorated by different types of modifications both in bacteria and mammal (6). In the 16S rRNA of bacteria (SSU rRNA), C1402 (corresponding to C1702 in human) is sequentially modified by two methyltransferases, rsmI and rsmH, to form m^4^Cm1402 (35), but the partially paired A1509 (A1832 in human) on the opposite side is without any modification (Fig. S5A, left panel) (6). The Cryo-EM structure of the bacterial ribosome shows direct interactions of the N^4^-methyl group of m^4^Cm1402 with the backbone of mRNA and is important for translation fidelity (35). Intriguingly, human 18S rRNA has a different modification pattern; i.e., the adenosine A1832, which corresponds to A1509 in bacteria, is instead modified by m^6^A while the opposite cytosine (C1702, which is m^4^Cm methylated in bacteria) is only modified by 2’-O methylation without further N^4^-methylation (Fig. S5A, middle panel). The underlying reasons for this transition from methylation of a cytosine in bacteria to adenine in humans are still unclear. Given the abundance of rRNAs, considerable amounts of m^4^C(m) can be generated from rRNA degradation, but unlike bacteria, metazoa and plants do not have a m^4^C(m) clearance system to remove the potentially toxic, modified m^4^C(m) base. This may be the selective pressure that drives the transition from m^4^C(m) to the metabolizable m^6^A in metazoan and plants. Consistently, mammalian and plant cells have evolved to have an N^6^-methyl-AMP deaminase (MAPDA) to protect cells from misincorporation of m^6^A resulting from RNA degradation into transcribing RNA (36). Similar to *METTL5*, the *MAPDA* gene is also mainly found in metazoa and plants (36,37).

### Do rRNA modifications play a role in ribosomal heterogeneity?

The small ribosomal subunit decodes genetic information within mRNA, whereas the large subunit covalently links amino acids into a nascent protein (38). As discussed earlier, the 18S m^6^A1832 is a conserved modification localized in the key region of the ribosome small subunit, and therefore its conservation suggests that it’s subject to selective pressure during evolution. Surprisingly, the 18S m^6^A1832 is not essential for ribosomal biogenesis and loss of methylation doesn’t have any detectable effects on global translation, which might be due to functional redundancy with other modifications that occur on rRNAs and compensation by changes of those modifications upon the loss of 18S m^6^A1832. An alternative possibility is that this “non-essential” modification may be leveraged to increase the diversity of ribosome, i.e., ribosomal heterogeneity (39–41), in order to cope with the increasingly complex nature of mRNA in higher species or functions as a sensor to fine-tune expression of a subset of mRNAs with specific roles in specific biological contexts. Although the latter is an attractive hypothesis, the observation that A1832 of the 18S rRNA is almost fully methylated would appear to be inconsistent with this model. However, it remains possible that dynamic regulation of rRNA methylation may be cell typespecific, and in cell lines where METTL5 plays a critical functional role (DepMap, 17/739) may display a more dynamic 18S rRNA methylation pattern. It will be important to determine if any cell types changes 18S rRNA methylation level in response to the ISR to understand if rRNA methylation *per se* plays a role in the ISR.

In addition to our findings, emerging published evidence also demonstrates that differential expression and post-translational modifications of ribosomal proteins might contribute to the assembly of function-specific translational machines. For instance, the presence of the testis-specific RPSY2 in the testis ribosome suggests tissue-specific ribosomal machines (42). Akin to METTL5, loss of the large subunit ribosomal protein Rpl38 doesn’t affect global protein synthesis, but perturbs translation of a select subset of Hox mRNAs (43). Additionally, a repertoire of 2’-O-methylation sites in rRNA were identified to be subjected to variations and it was demonstrated that functional domains of ribosomes are targets of 2’-O-methylation plasticity (44). Moreover, the 28S rRNA A4220 is not always 100% methylated, e.g., it is about 50% level in HepG2 cells (9), suggesting that HepG2 cells may have ribosomes with and without m^6^A4220. Collectively, these findings argue that rRNA modifications may contribute to the formation of ribosomes with different configurations, thus contributing to ribosomal heterogeneity. However, exactly how these “specialized” ribosomes selectively impact translation of only a small subset of mRNAs is unclear and it will be interesting and important to investigate the underlying molecular mechanism that govern this selectivity in the future.

### METTL5, ISR and tumorigenesis?

The ISR is characterized by the phosphorylation of eIF2α and activation of ATF4 translation in response to a variety of stress signals, including ER stress (17). Our results show that Mettl5 loss does not affect eIF2α protein level and its phosphorylation. Instead, Mettl5-mediated 18S rRNA methylation appears to be important for translational control of Atf4, suggesting that Mettl5 does not work in the classical senor kinases-eIF2α-Atf4 pathway. Indeed, blocking sensor kinase activities by the small molecule inhibitor GSK2606414 (17) caused further decrease of Atf4 target gene transcription in Mettl5 KO cells, supporting the hypothesis that Mettl5 likely works in parallel to the known ISR pathway.

The transcription factor ATF4 plays a central role in response to unfavorable conditions such as accumulation of unfolded or mis-folded proteins, nutrient depletion and DNA damage. Tumor cells inevitably encounter various types of stress, including hypoxia, limited glucose and amino acids shortage (14,45). ATF4 is often found to be up-regulated in tumor cells to promote adaptation of cells to these challenges (46). Interestingly, the METTL5 expression level is noticeably up-regulated in tumors, especially solid tumors, compared with the normal control tissues in various TCGA tumor datasets (Fig. S5B). Importantly, the expression of the METTL5 cofactor, TRMT112, is also up-regulated and correlates well with the expression pattern of METTL5 in these tumors (Fig. S5C). We speculate that up-regulation of METTL5 may serve to ensure 18S rRNA methylation in order to sustain a high level of ATF4 protein expression (through regulation of translation efficiency of ATF4) to counter the stress in tumors. It should be noted that our data show that METTL5 is necessary for productive ATF4 translation even in the absence of any stress inducers suggesting that METTL5, and hence 18S rRNA methylation, is necessary for basal translation of ATF4.

In summary, we identify METTL5-TRMT112 as an m^6^A enzymatic complex that specifically mediates m^6^A methylation of 18S rRNA at residue A1832, and this modification is conserved and critical for translation of a subset of mRNAs. METTL5 depletion leads to inefficient translation of stress-responsive genes including ATF4, and compromises ISR (Fig. 6). Our findings highlight METTL5 and possibly rRNA methylation as a mechanism to fine-tune mRNA translation, possibly by contributing to the formation of specialized ribosomes and/or the proper conformation of ribosomes that favors the optimal translation of selective mRNAs. Our data, however, do not exclude the less likely possibility that METTL5 may regulate expression of the stress-related gene indirectly or via a different, yet-to-be-identified substrate or METTL5 functions in an enzymatic activity independent manner. Finally, the fact that METTL5 is frequently up-regulated in various tumor types raises the exciting possibility that METTL5 may be a promising target for tumor therapy in the future.

**Figure 6.**
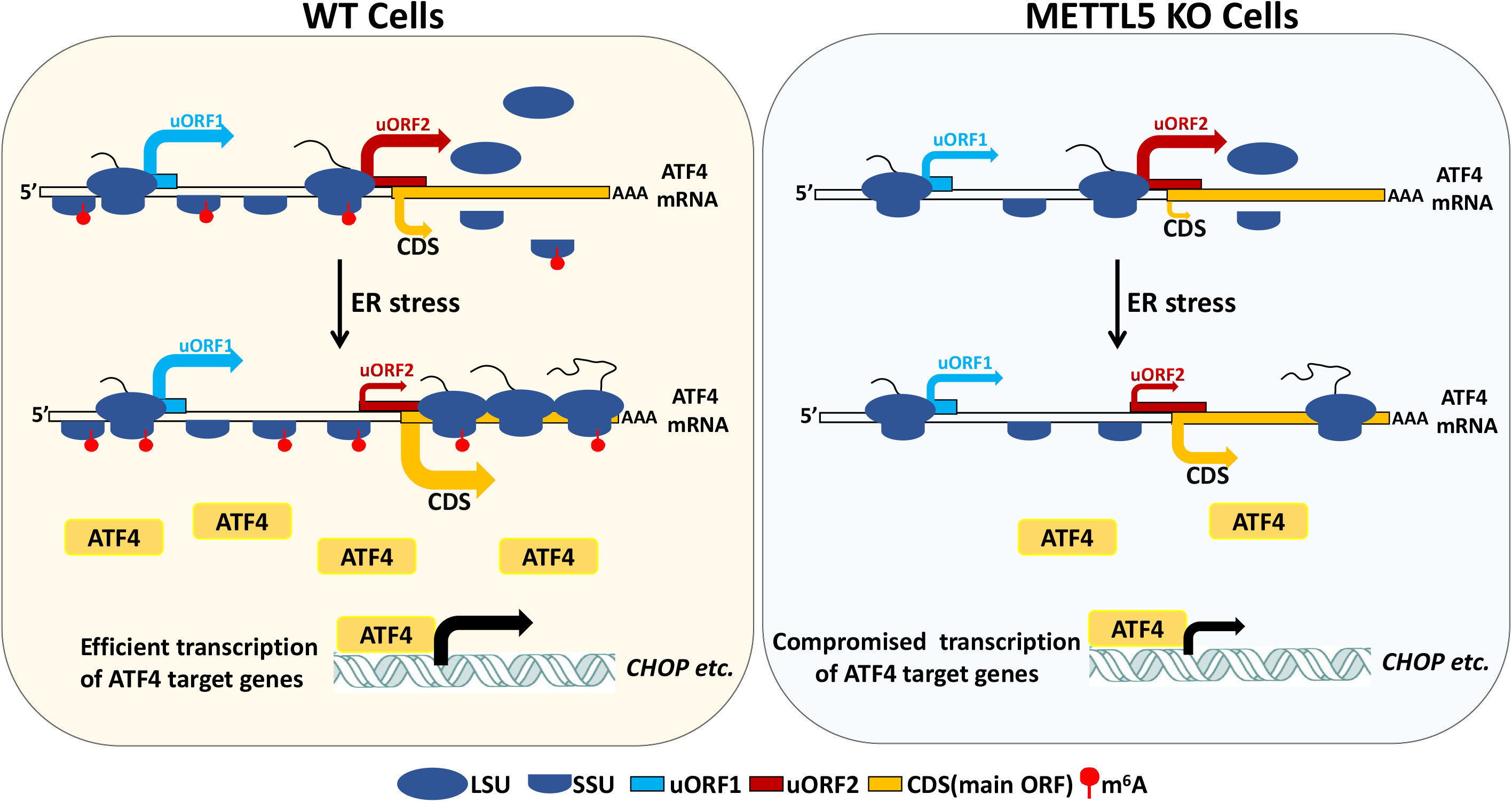
A working model for METTL5 regulating ATF4 expression. METTL5 modified 18S rRNA is critical for maintaining efficient translation of *ATF4*. In the METTL5 KO cells (right: light blue box), basal translation of uORF1 and CDS is greatly hindered, presumably to be due to ribosome conformation changes caused by the lack of m^6^A modification in the 18S rRNA. Upon ER stress, *ATF4* mRNA is efficiently translated by the m^6^A-decorated ribosome in WT cells, followed by acute up-regulation of a set of its downstream targets (*CHOP etc.*). However, the activated translation of *ATF4* is compromised when 18S rRNA m^6^A methylation is absent, as the unmodified ribosome seems less favorable to going through translation initiating from uORF1 and the main ORF (CDS), thus reducing productive translation of sufficient ATF4 proteins in response to stress.

## Experimental Procedures

### Plasmids and protein purification

For the expression of METTL5 protein, METTL5 ORF cDNA was cloned into pHAGE or pET28a expression vector (Invitrogen), respectively. METTL5 mutants were generated using the QuikChange Site-Directed Mutagenesis Kit (Stratagene) according to the manufacturer’s protocol. The shRNAs targeting different methyltransferases were constructed into pLKO.1 vector. METTL5-specific 20 nt guide RNA sequences were cloned into pLenti-CRISPR v2 plasmid.

Recombinant METTL5 was expressed in the Rosetta (DE3) bacterial cells, which were incubated at 37 °C until OD600 reached ~ 0.6–1 and then cooled down to 16 °C. IPTG was added to 0.2 mM final concentration, and cells were further incubated at 16 °C for 16 h. Cell pellets were lysed in a buffer containing 300 mM NaCl, 25 mM Tris pH 7.5, 10% glycerol, 0.5% NP-40. Total lysate was incubated with HisPur Ni-NTA Resin at 4 degree for 5 hours. His-METTL5 protein was eluted with elution buffer (20 mM Tris pH7.5, 150 mM NaCl, 200 mM Imidazole) in 0.5 ml aliquots until color change was no longer observed (Bradford assay).

For purification of the METTL5/TRMT112 complex, METTL5 was cloned into pET28b vector with 6xHis-SUMO affinity tag (pXC2097), and TRMT112 was cloned in pET22b vector without any tag (pXC2076), and then two plasmids were co-transformed into *Escherichia coli* Rosetta cells. Coexpression of METTL5 and TRMT112 was performed at 16 °C overnight after induction with 0.1 mM IPTG. Cells were harvested and sonicated in lysis buffer of 20 mM Tris-HCl, pH 8.0, 150 mM NaCl, 5% glycerol, 0.5 mM tris (2-carboxyethyl) phosphine (TCEP). The supernatant was loaded onto a Ni-NTA affinity column and washed with lysis buffer containing 50 mM imidazole. The METTL5/TRMT112 complex was eluted with buffer containing 250 mM imidazole. After removing the N-terminal His_6_-SUMO tag by Ulp1 protease (purified in-house) overnight at 4°C, the target complex was further purified by ion-exchange chromatography (HiTrap Q-HP, GE Healthcare), and finally purified by size-exclusion chromatography (Superdex 200, GE Healthcare) in 20 mM Tris-HCl (pH 8.0), 150 mM NaCl, 5% glycerol, 0.5 mM TCEP. All purification steps were performed at 4°C.

### Assays for m^6^A methyltransferase activity *in vitro*

*In vitro* methyltransferase activity assays were performed in a 30 μL reaction mixture containing the following components: 1 μg RNA probes, 0.5 μg fresh recombinant protein, 0.8 mM *d3*-SAM, 50 mM Tris–HCl, 5 mM MgCl_2_, 1 mM DTT, pH 8.0. The reactions were incubated at 37 °C for 1hr. After incubation, samples were treated with proteinase K at 50 °C for 20 min, and the resultant RNA was desalted and then digested with nuclease P1 and alkaline phosphatase. Nucleosides were quantified by using nucleoside-to-base ion mass transitions of 285 to 153 (*d3*-m^6^A) and 268 to 136 (Adenosine). Adenosine (A) served as an internal control to calculate the amount of RNA probe in each reaction mixture for QQQ LC–MS/MS analysis.

### Steady-state kinetic measurement

Single-strand 14-nt RNA corresponding to the sequence surrounding A1832 of 18S rRNA (5’-UCGUA**A**CAAGGUUU-3’) was used as a substrate for the METTL5/TRMT112 complex. Reactions were carried out in 20 μl mixtures containing 20 mM Tris-HCl (pH 8.0), 1 mM DTT and 0.2 μM METTL5/TRMT112 complex. The reactions were allowed to proceed at room temperature for indicated durations (5, 10, 15 and 20 min), and were terminated by the addition of TFA (Trifluoroacetic acid) to a final concentration of 0.1% (v/v). Then, 5 μl of reaction mixture was transferred to a low-volume 384-well plate, and the methylation activity was measured using the Promega bioluminescence assay (MTase-Glo™) (47). The bioluminescence assay was performed according to the manufacturer’s protocol, in which the by-product SAH of the methylation reaction is converted into ATP in a two-step reaction and ATP is detected through a luciferase reaction. The luminescence signal was measured using a Synergy 4 Multi-Mode Microplate Reader (BioTek).

### Antibodies and Chemical reagents

Anti-FLAG (M2) beads and antibody (F1804) were purchased from Sigma. Anti-HA beads (sc-7392ac) were anti-HA antibody (#2367S) were provided/purchased by/from Santa Cruz and CST, respectively. Anti-METTL5 (16791-1-AP) and anti-ATF4 (10835-1-AP) antibodies were purchased from Proteintech. Anti-TRMT112 (sc-398481) antibody was provided by Santa Cruz.

Chemical activators/inhibitors used in this study include: GSK2606414 (PERK inhibitor, 17376, Cayman Chemical), Thapsigargin (Unfolded protein response inducer, 67526-95-8,| Cayman Chemical) and Cycloheximide (Protein synthesis inhibitor, Sigma, C7698).

### Cell Culture, Transfection and Inhibitor Treatment

Cells were cultured in DMEM supplemented with 10% fetal bovine serum (FBS) (Hyclone) and 1% Pen-Strep unless otherwise specified. Cell transfection was performed using Lipofectamine 2000 (Invitrogen). For the inhibitor treatment experiments in general, cells were treated with 0.5μM GSK2606414 (GSK) for 1 hours; 2μM Thapsigargin (TG) for 3 hours; Cycloheximide (CHX) was used at the final concentration of 100 μM.

### Co-immunoprecipitation

Cells were lysed with the lysis buffer (20 mM TrisCl (pH 7.5), 150 mM NaCl, 1 mM EDTA, 0.5% NP-40, protease inhibitors and phosphatase inhibitor cocktail (Roche Applied Science). The extract was spun at 14,000 rpm for 15 min at 4°C. Two percent of the extract was kept as input, while the rest was incubated with the appropriate antibody (1-2 μg) for 1 h at 4°C. Protein A/G beads (Millipore) were then added for overnight incubation at 4°C. The beads were washed five times with the lysis buffer, and the bound proteins were boiled in SDS sample buffer and Western blotted using indicated antibodies.

### Isolation of full-length rRNA and rRNA fragment for QQQ (Triple Quadrupole) LC-MS/MS analysis

To purify the 18S and 28S rRNAs, total RNA was separated by electrophoresis with 1% Native TAE agarose gel. 18S and 28S RNAs were extracted from gel slices using the Zymoclean Gel RNA Recovery kit for HPLC-MS/MS analysis. To isolate the region of 18S rRNA that contains A1832, 2000 pmol of a synthetic oligo deoxynucleotide (Biotin-acctacggaaaccttgttacgacttttacttcctct, synthesized in (Integrated IDT DNA Technologies, Inc.)) complementary to the A1832 surrounding region of 18S rRNA were incubated with 200 pmol of total RNA, in 0.3 volumes of the hybridization buffer (250 mM HEPES pH 7, 500 mM KCl). The hybridization mixture was incubated at 90 °C for 7 min and slowly cooled to room temperature (25 °C) over 3.5 h to allow hybridization to occur. Single-stranded RNA and DNA were digested with mung bean nuclease (New England BioLabs) and RNase A over 1 h at 37 °C in a buffer solution (50 mM NaOAc pH 5, 30 mM NaCl, and 2.5 mM ZnCl_2_). Digestion products were mixed with streptavidin T1 beads in IP buffer (150 mM NaCl, 50 mM Tris, pH7.9, 0.1% NP-40) at 4 °C for 1 h. After washing with IP buffer, beads were heated at 70 °C for 5 min and supernatant (containing 40 nt RNA fragment) was collected.

### QQQ LC–MS/MS

200 ng of RNA were digested by nuclease P1 (1 U, Sigma) in 25 μl of buffer containing 25 mM NaCl and 2.5 mM ZnCl_2_ at 42 °C for 2 h, which was followed by the addition of NH_4_HCO_3_ (1 M, 3 μl, freshly made) and Calf Intestinal Alkaline Phosphatase (1 U, Sigma) and additional incubation at 37 °C for 2 h. Samples were then diluted to 60 μl and filtered (0.22 μm pore size, 4 mm diameter, Millipore); 5 μl of solution was loaded into LC–MS/ MS (Agilent6410 QQQ triple-quadrupole mass spectrometer). Nucleosides were quantified by using retention time and nucleoside to base ion mass transitions of 282.1 to 150.1 (m^6^A), and 268 to 136 (A).

### Enhanced-CLIP (eCLIP) and RNA sequencing

eCLIP was carried out according to the previously reported procedures (32) using 293T cell lines stably expressing HA-tagged METTL5. Briefly, cells were irradiated on ice, using UV crosslinking (254 nm, 400 mJ/cm^2^). HA-METTL5 was immunoprecipitated with anti-HA magnetic beads. Then, on-bead digestion was performed with RNase I (Ambion), and stringent washes were carried out 4 times. The bound RNAs were extracted from SDS–PAGE gel, and RNA libraries were prepared as described (32). Qualified RNA libraries were sequenced using Illumina NovaSeq with pair-end 150-bp read length. Each experiment was conducted with two biological replicates and analyzed as described (32,48).

### Polysome profiling

Polysome profiling was adapted from the previously reported procedure (34). Two 10-cm plates of 293T cells were prepared for each sample (wild-type and METTL5 KO). Cells were treated with cycloheximide (CHX), at 100 μg/ml for 10 min before collection. Cells were lysed on ice for 30 min in 1 ml lysis buffer (10 mM Tris–HCl pH7.4, 5 mM MgCl_2_, 100 mM KCl, 1% Triton X-100, 2 mM DTT, 500 U/ml RNase inhibitor, 100 μg/ml CHX, protease inhibitor cocktail). Supernatant (~1 ml) was collected, and absorbance at 260 nm (A260) was measured. Samples with equal A260 readings were loaded onto a 10–50% (w/v) sucrose gradient buffer (20 mM HEPES, pH 7.4, 100 mM KCl, 5 mM MgCl_2_, 2 mM DTT, 100 μg/ml cycloheximide, 1 × protease inhibitor cocktail (EDTA-free), 20 U/ml SUPERase inhibitor) prepared using gradient station (Biocomp). Gradients were centrifuged at 4 °C for 4 h at 28,000 r.p.m. (Beckman, rotor SW28). Samples were then fractioned and analyzed by Gradient Station (Biocomp) equipped with an ECONO UV monitor (Bio-Rad) and fraction collector (FC203B, Gilson).

### Ribo-seq

For Ribo-seq, mouse B16 cells were pre-treated with 50 μg/ml of cycloheximide (CHX) for 10 min, and then washed twice with ice cold PBS supplemented with CHX. Cells were collected and re-suspended in 400 μL lysis buffer (20 mM Tris-HCl pH 7.4, 150 mM NaCl, 5 mM MgCl_2_, 1 mM DTT and 100 mg/ml CHX). After incubation on ice for 10 min, lysate was triturated 5 times through a 25-gauge needle and then lysate was centrifuged at 20,000 x g for 10 min. 5 μL of lysate was flash frozen and saved as input. To generate ribosome-protected fragments the lysates (30 mg) were first mixed with 200 μL DEPC-H2O then incubated with 15 U RNase I for 45 min at room temperature. The reaction was stopped with 10 μL SUPERase*In RNase inhibitor. 0.9 mL of sucrose-supplemented lysis buffer was added to the digestion mixture and ultracentrifuged at 100,000 rpm (TLA100.3 rotor), 4 °C for 1 h. Pellets were re-suspended in 300 μL of water and after phenol-chloroform extraction, precipitated with ethanol. The RNA was then run on a 15% 8 M urea TBE gel, stained with SYBR Gold, and a gel fragment between 17-34 nucleotides corresponding to ribosome-protected RNA was excised. RNA was eluted for 2 h at 37 °C in 300 μL RNA extraction buffer (300 mM NaOAc pH 5.5, 1 mM EDTA, 0.25%v/v SDS) after crushing the gel fragment. RNA was ethanol precipitated and re-suspended in 26 μL water and treated with RiboZero Gold kit. Purified RNA was treated with PNK and subjected to library construction with NEBNext Small RNA Library Prep Kit.

### Ribo-Seq data analysis

The clean reads were obtained by trimming the adapters and filtering the low-quality sequences (Q30) from raw FASTQ sequences of input and ribosome profiling using trim_galore (https://www.bioinformatics.babraham.ac.uk/projects/trim_galore). For input samples, the clean reads were then mapped to mouse reference genome (mm10) using STAR (49). The mapping results in BAM format were used as the input of featureCounts program from the Subread package (50) to calculate the raw read count for CDS region of each gene in GENCODE mouse gene mode (M21). For the clean Ribo-seq data, the clean ribosome-protected fragments (RPFs) were collapsed into FASTA format by fq2collapedFa.pl (51). Ribo-toolkit (https://bioinformatics.caf.ac.cn/) was used to perform various analyses based on the collapsed RPF tags and raw read counts from input samples, which include tRNA and rRNA removal, RPF quality controls, codon occupancy analysis and translation efficiency analysis. In brief, the RPFs tags were first aligned to rRNA sequences from Ensembl database (release 91) and tRNA sequences from GtRNAdb databases (52) to exclude the tags derived from rRNAs and tRNAs. The filtered RPF tags were then mapped to mouse reference genome using STAR (49) and the count of unique-mapped RPF tags on CDS region of each GENCODE gene mode (M21) was computed using featureCounts. The translation efficiency (TE) was calculated by dividing normalized RPF abundance by its normalized mRNA abundance. A threshold of ΔTE (fold change of TE) > ±1.5 and FDR <0.05 was used to define the differentially translated genes (53). For codon occupancy analysis, the genome unique mapped tags were aligned against mouse transcript sequences (GENCODE M21) using Bowtie with a maximum one mismatch allowed (54). Then, an offset of 15 nt from 5’ end of 27-32nt RPFs translated in the zero-frame of CDS were used to infer the A site codon occupancy, which was further normalized by its basal occupancy (the average codon occupancy among +1, +2 and +3 downstream of A-site).

### Computation of MFE

MFE was computed using two programs: RNAfold, included in the ViennaRNA software package version 2.1.5; and Quickfold, from the Mfold web server (http://mfold.rna.albany.edu/?q=DINAMelt/Quickfold). For very short sequences, we found that the MFEs computed by Quickfold (Mfold) were sometimes positive. In these cases, global free energy were set to 0 kcal/mol.

### Gene Ontology and Gene Set Enrichment Analysis

Gene Ontology analysis of differentially translated genes was conducted using DAVID GO (55). To identify the enriched ontology terms, a significance threshold based on Benjamini-Hochberg adjusted p-value (FDR) < 0.05 was used. For Gene Set Enrichment Analysis (GSEA) (56), the Java implementation from Broad institute (http://software.broadinstitute.org/gsea/index.jsp) was used to evaluate the statistical enrichment of gene ontology sets in the MSigDB C5 gene ontology collection. Briefly, the translation gene list containing 10,446 genes was pre-ranked by fold change, which was then used as the input of the Java implementation to calculate the normalized enrichments score and adjust p-values based on classic mode.

### TCGA data analysis

RNA-Seq expression data and small RNA-Seq data for 33 TCGA tumor types were downloaded from Genomic Data Commons Data Portal (GDC) of TCGA (http://cancergenome.nih.gov/). The gene expression matrix was then constructed by merging the TPM (Transcripts Per Million) values of all RNA-seq samples. The expression correlations between METTL5 and TRMT112 among TCGA tumors were conducted by using Pearson correlation coefficient.

## Acknowledgments

We thank Erdem Sendinc and Bill Neidermyer for assistance with setting up the ribosome profiling experiments and their insightful advice. We also thank Santa Cruz Biotechnology, Inc. for providing antibodies. This work was supported by an R35 grant from the NCI (1R35CA210104) and by funds from BCH to YS. This work was also supported in part by grants from the NIH to X.C. (R01 GM049245), and a CPRIT grant to X.C. (RR160029).

## Conflict of interest

Y.S. is a co-founder and equity holder of Constellation Pharmaceuticals and Athelas Therapeutics, a consultant for Active Motif, Inc and has equity in Imago Biosciences. All other authors declare no competing financial interests.

## Author contributions

Y.S. and H.C. conceived and designed the project. H.C., C.Z. and H.C. carried out most of the biochemical and cellular experiments. Q.L. performed the bioinformatic analysis. P.H. was responsible for mass spectrometry analysis. D.Y. performed protein purification and kinetic experiments under the supervision of X.Z. and X.C. B.K. and R.G. provided advice on the possible role of 18S rRNA methylation on ribosome conformation and Ribo-seq data analysis, respectively. Y.S. supervised the project throughout. Y.S. and H.C. cowrote the manuscript and all authors contributed to the manuscript writing.

## Data Availability

Sequence data from this study have been deposited in the Gene Expression Omnibus with accession code Ribosome profiling: GSE138110.

## Footnotes

1. This study was supported by Boston Chilren’s HospitalBCH funds to YS; 2. Y.S. is also an American Cancer Society Research Professor; 3. During the preparation of this manuscript for publication, two other studies have been published that also documented the role of METTL5 as an 18S rRNA-specific methyltransferase (57,58). And in one of the studies METTL5 has also been shown to regulate stress-related genes in *C. elegans* (58), which suggests the involvement of METTL5 in stress response may be conserved during evolution.

---

**sFigure 1.**
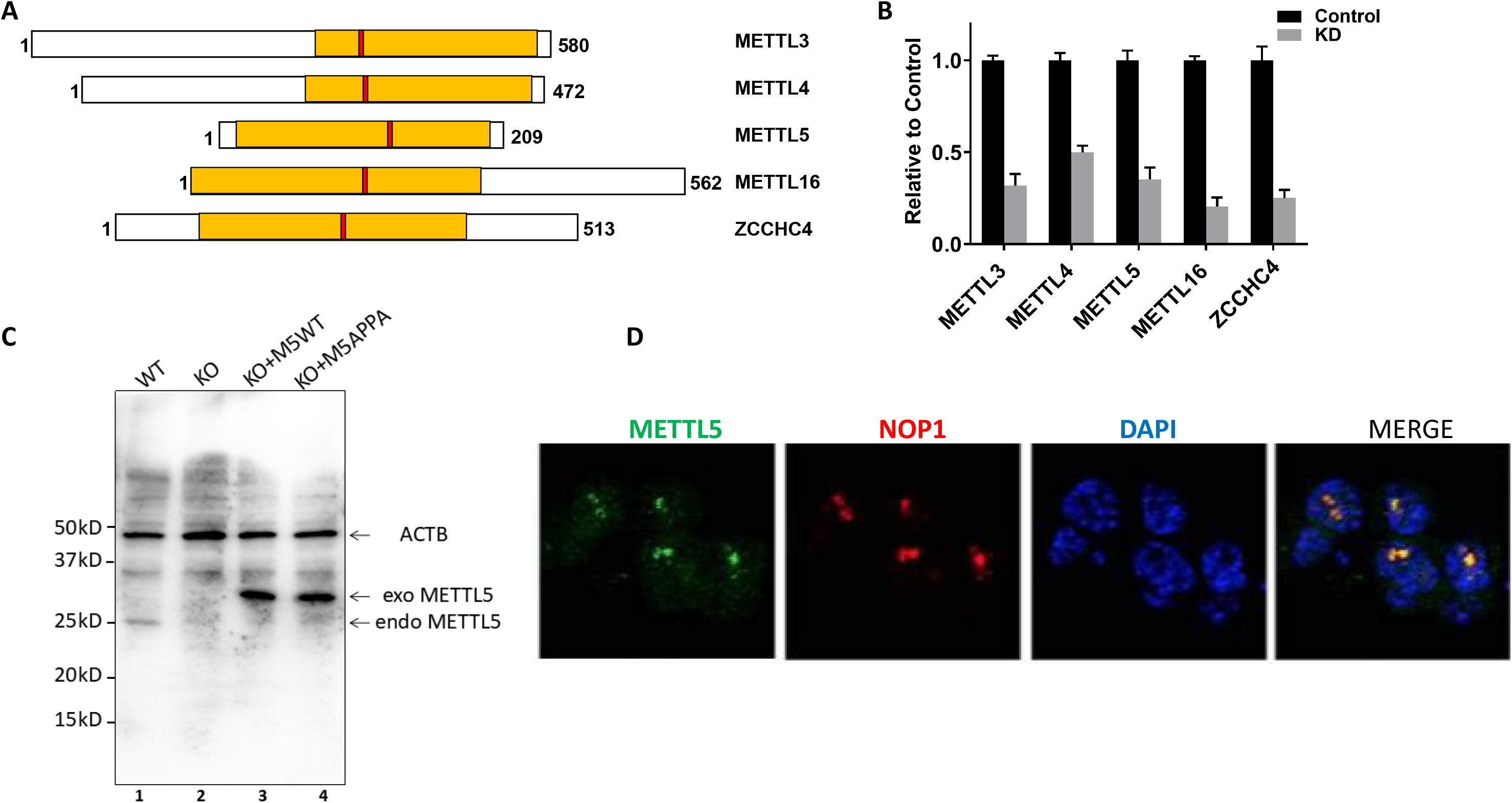
A. Schematic diagram of the putative m6A methyltransferase domain in METTL family proteins, yellow: methyltransferase domain, red: [DNSH]PP[YFW] motif; B. RT-qPCR verification of KD efficiency for the potential methyltransferases; C. Verification of KO and rescue of METTL5 in 293T cells by WB. The cell lysates from indicated stable cell lines were blotted with METTL5 and β-actin (ACTB) antibodies was included as a loading control, Exo METTL5: exogenously expressed METTL5 protein tagged with HA epitope, endo METTL5: endogenous METTL5; D. The cellular distribution of METTL5 was shown to be mainly in nucleoli by fluorescence microscopy through immunofluorescence staining with METTL5 antibodies. NOP1, a well-characterized nucleoli protein, was used to mark nucleoli. Co-localization of the endogenous METTL5 with NOP-1 is shown.

**sFigure 2.**
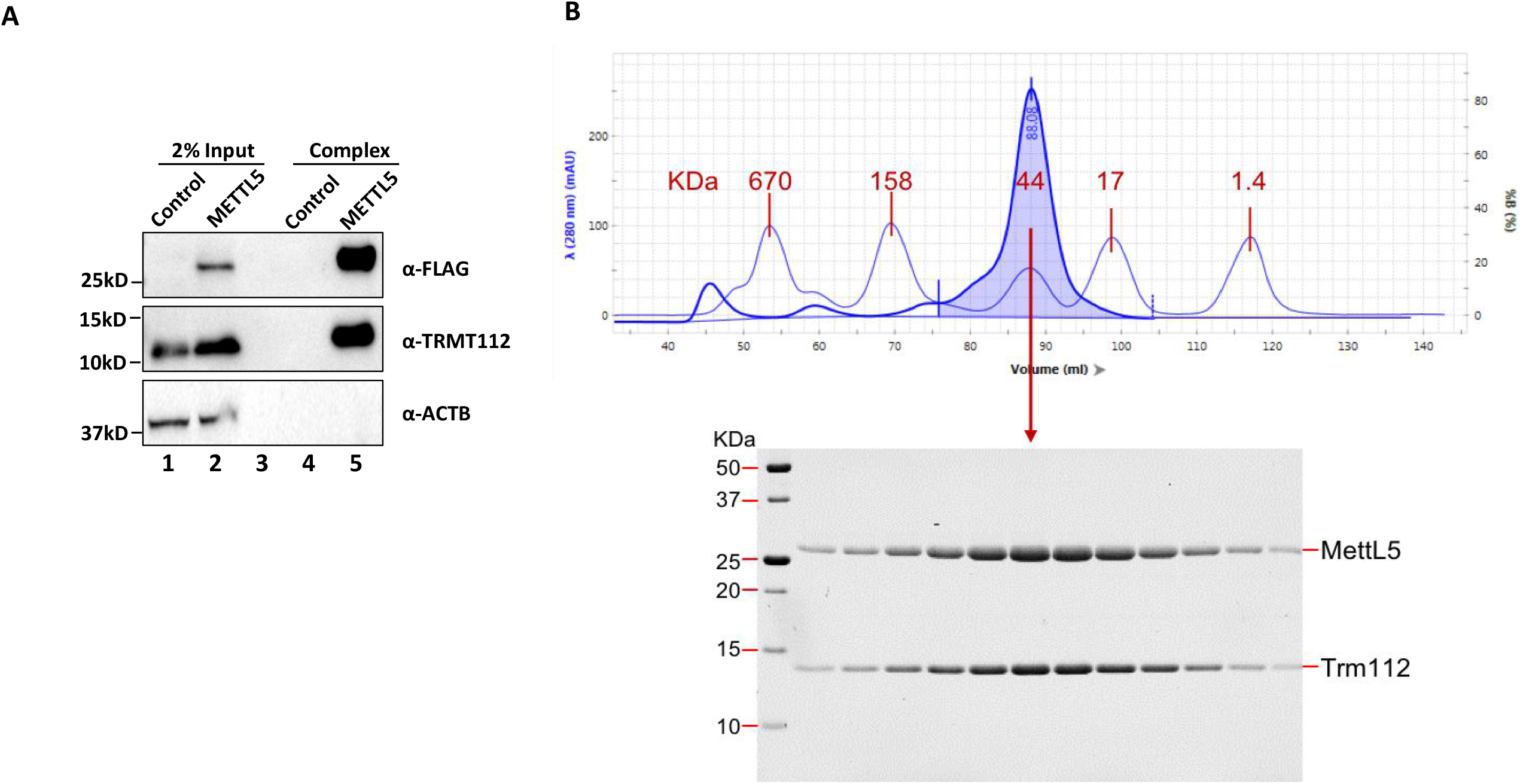
A. A list of proteins identified by LC-MS/MS in the METTL5 pulldown. TRMT112 was highlighted with red bold fonts; B. WB using TRMT112 antibodies readily detected the existence of TRMT112 in the purified METTL5 complex, which independently validated the MS/MS results; C. METTL5 and TRMT112 form a complex and were co-purified via a 4-column chromatography. (top) Profile of size-exclusion chromatography (Superdex 200), superimposed with standards indicated by molecular weight in kDa; (bottom) Bio-Rad 18% Stain-free PAGE of peak fractions.

**sFigure 3.**
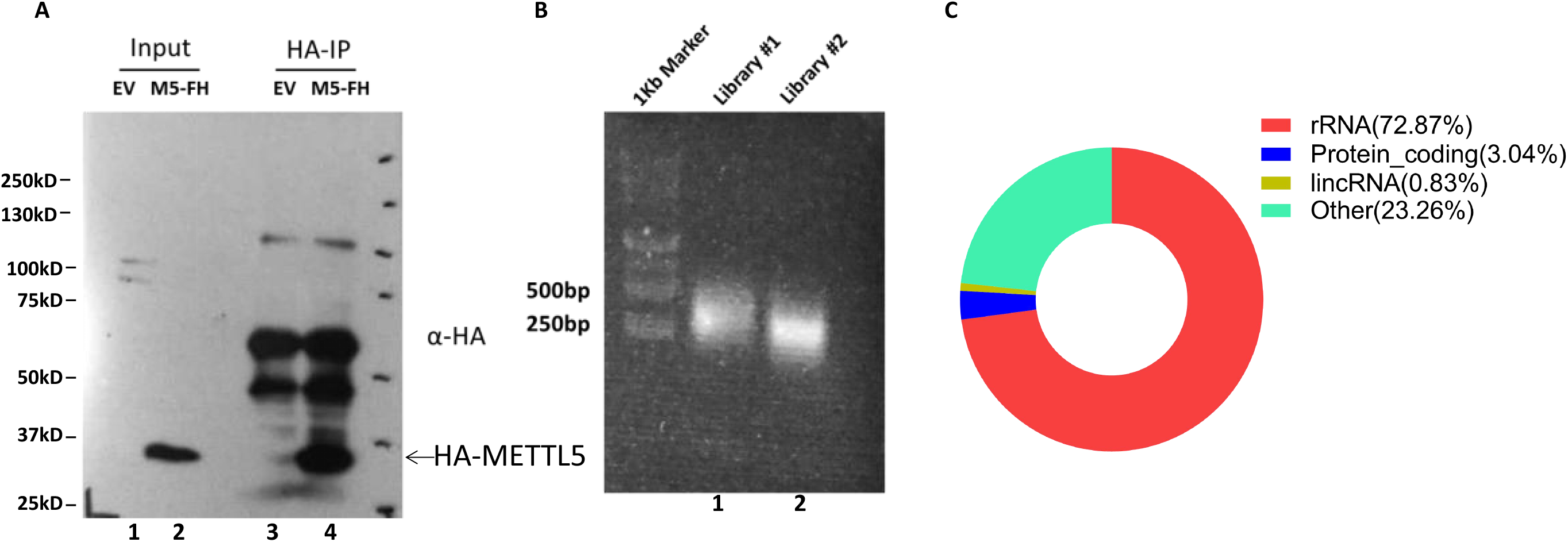
A & B. The preparation of METTL5-HA eCLIP libraries. The RNAs bound with METTL5 were recovered from Nitrocellulose membrane and subjected to library construction as described in the Methods; C. The identified peaks in eCLIP experiment were subdivided into four categories based on the RNA types, among which rRNA is the dominant type (>70%) among RNAs that are bound by METTL5.

**sFigure 4.**
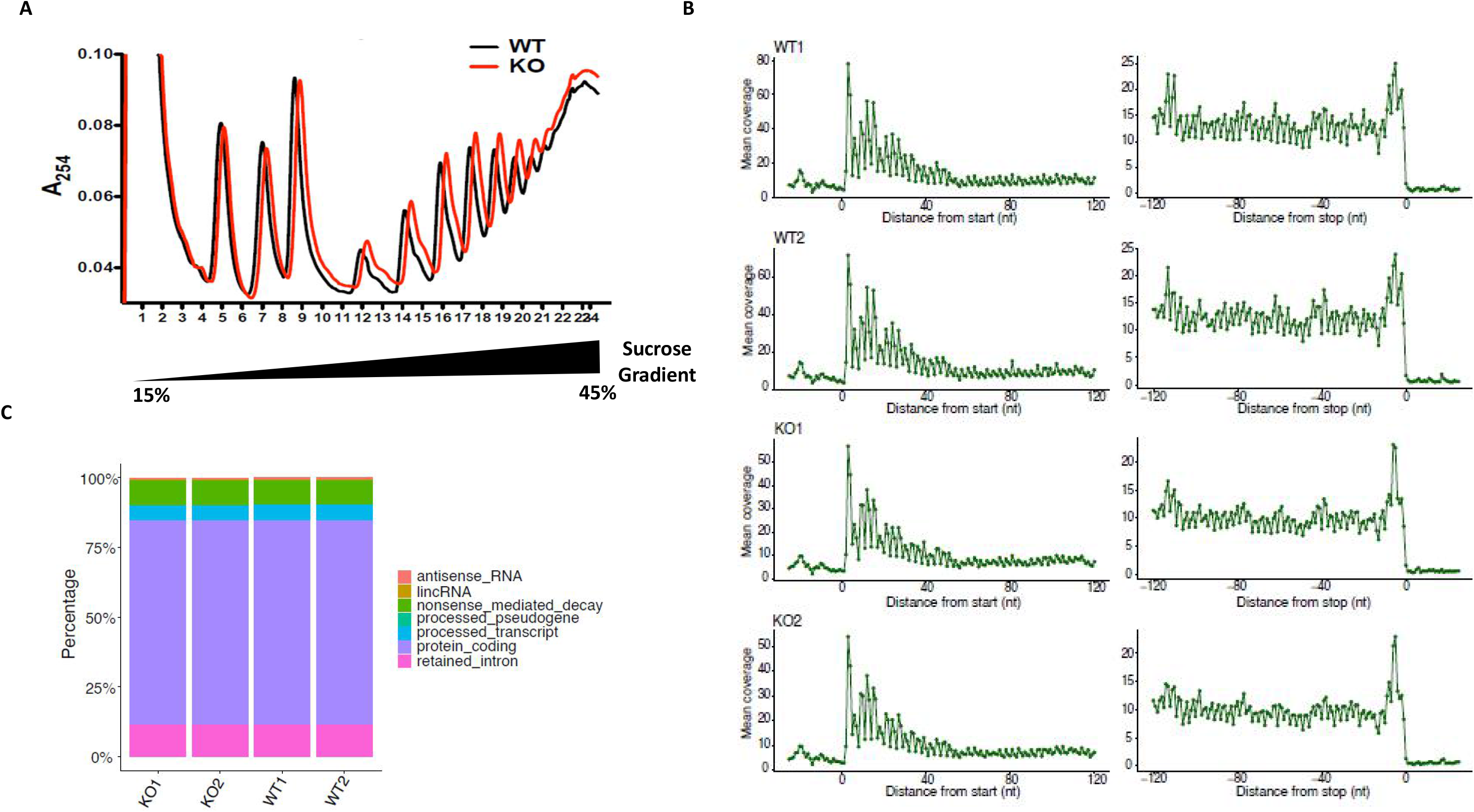
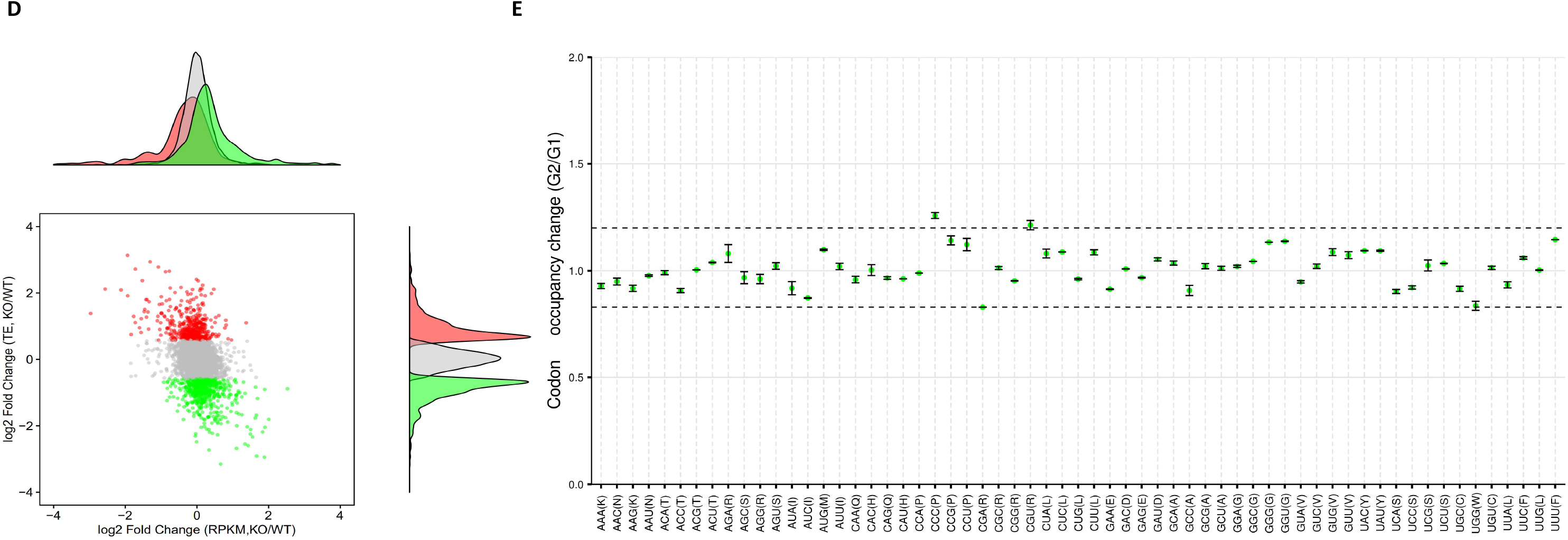
A.Ribosomal subunit profiling by sucrose density gradient under low Mg2+ concentration (0.5 mM). There were no detectable differences between WT and KO 293T cells, which indicated that METTL5 KO doesn’t affect ribosome assembly or global translation. n=3 independent experiments; B. Meta-gene analysis of RPF fragments distribution around Translation Starting Site (TSS) and Translation Ending Site (TES). C. The relative counts of RPF fragments in different types of RNAs; D. The pattern of translation and RNA level changes of expressed genes in WT and KO B16 cells (grey: TE unaffected, green: TE down-regulated, red: TE up-regulated). The grey, green and red peaks are well overlapped, which indicate that the observed translational efficiency differences are not caused by changes in gene transcription; E. There are no significant differences in codon (except CCC and CGU) usage between WT and KO B16 cells.

**sFigure 5.**
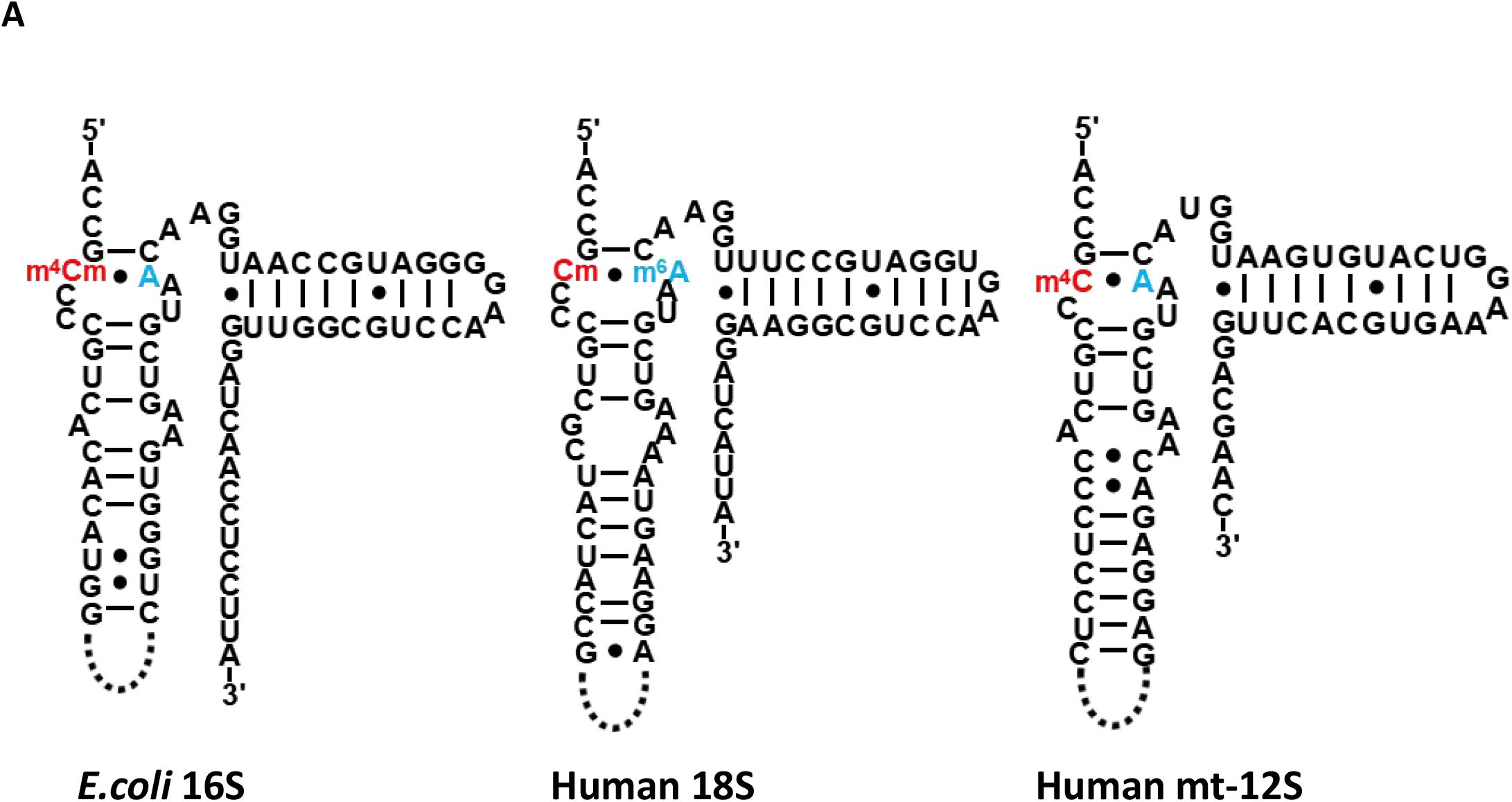
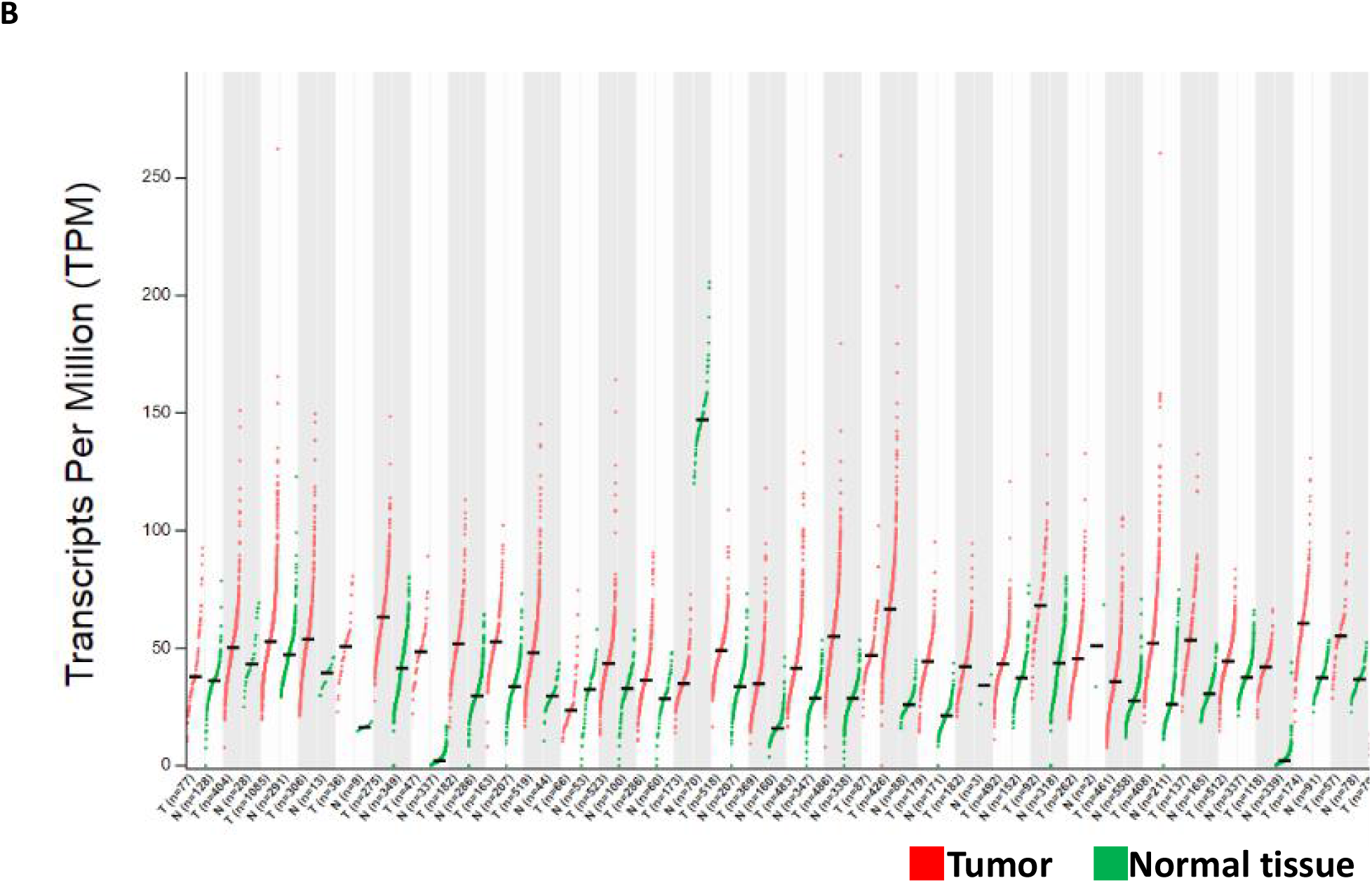
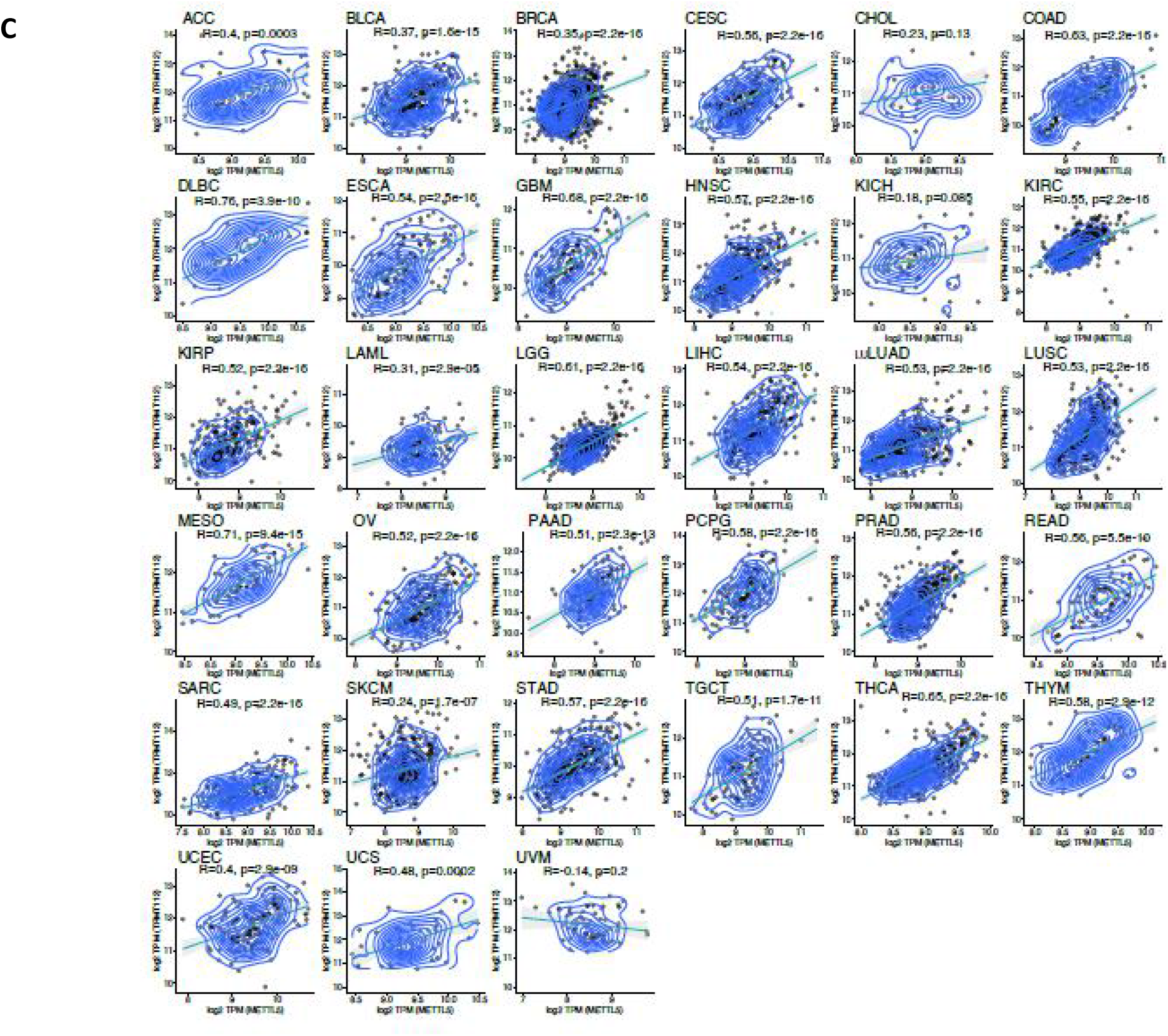
A. SSU rRNA methylation pattern changes during evolution. The 3’-terminal region is decorated by distinct types of modifications in different SSU rRNAs. In E.coli 16S and human mt-12S rRNA, the cytosine is modified by N4-methylation, which is then paired with an unmodified adenosine. In contrast, the equivalent adenosine is methylated in human (and other animals or plants) 18S rRNA, and is paired with an unmodified cytosine; B. METTL5 is up-regulated in various tumors. The gene expression data-sets were retrieved from TCGA and the differences in various tumor and normal tissues were compared using GEPIA2 software(http://gepia2.cancer-pku.cn/). Red: tumor, green: normal tissue; C. Expression of TRMT112 is up-regulated in various tumors and correlates well with METTL5 expression. The gene expression data of TRMT112 were also downloaded from TCGA and the correlation of these two genes’ expression was calculated in individual patients.

## Notes

### Competing Interest Statement

Y.S. is a co-founder and equity holder of Constellation Pharmaceuticals, Inc, a consultant for Guangzhou BeBetter Medicine Technology Co., LTD and Active Motif, Inc and an equity holder of Imago Biosciences. All other authors declare no competing financial interests.

### Summary of Updates

Acknowledgments and COI updated.

## References

1. Boccaletto, P., Machnicka, M. A., Purta, E., Piatkowski, P., Baginski, B., Wirecki, T. K., de Crecy-Lagard, V., Ross, R., Limbach, P. A., Kotter, A., Helm, M., and Bujnicki, J. M. (2018) MODOMICS: a database of RNA modification pathways. 2017 update. Nucleic acids research 46, D303–D307

2. Shi, H., Wei, J., and He, C. (2019) Where, When, and How: Context-Dependent Functions of RNA Methylation Writers, Readers, and Erasers. Molecular cell 74, 640–650

3. Meyer, K. D., and Jaffrey, S. R. (2017) Rethinking m(6)A Readers, Writers, and Erasers. Annual review of cell and developmental biology 33, 319–342

4. Yang, Y., Hsu, P. J., Chen, Y. S., and Yang, Y. G. (2018) Dynamic transcriptomic m(6)A decoration: writers, erasers, readers and functions in RNA metabolism. Cell research 28, 616–624

5. Natchiar, S. K., Myasnikov, A. G., Kratzat, H., Hazemann, I., and Klaholz, B. P. (2017) Visualization of chemical modifications in the human 80S ribosome structure. Nature 551, 472–477

6. Sergiev, P. V., Aleksashin, N. A., Chugunova, A. A., Polikanov, Y. S., and Dontsova, O. A. (2018) Structural and evolutionary insights into ribosomal RNA methylation. Nature chemical biology 14, 226–235

7. Sloan, K. E., Warda, A. S., Sharma, S., Entian, K. D., Lafontaine, D. L. J., and Bohnsack, M. T. (2017) Tuning the ribosome: The influence of rRNA modification on eukaryotic ribosome biogenesis and function. RNA biology 14, 1138–1152

8. Decatur, W. A., and Fournier, M. J. (2002) rRNA modifications and ribosome function. Trends in biochemical sciences 27, 344–351

9. Ma, H., Wang, X., Cai, J., Dai, Q., Natchiar, S. K., Lv, R., Chen, K., Lu, Z., Chen, H., Shi, Y. G., Lan, F., Fan, J., Klaholz, B. P., Pan, T., Shi, Y., and He, C. (2019) N(6-)Methyladenosine methyltransferase ZCCHC4 mediates ribosomal RNA methylation. Nature chemical biology 15, 88–94

10. Liu, B., and Qian, S. B. (2014) Translational reprogramming in cellular stress response. Wiley interdisciplinary reviews. RNA 5, 301–315

11. Spriggs, K. A., Bushell, M., and Willis, A. E. (2010) Translational regulation of gene expression during conditions of cell stress. Molecular cell 40, 228–237

12. Jackson, R. J., Hellen, C. U., and Pestova, T. V. (2010) The mechanism of eukaryotic translation initiation and principles of its regulation. Nature reviews. Molecular cell biology 11, 113–127

13. Hershey, J. W., Sonenberg, N., and Mathews, M. B. (2012) Principles of translational control: an overview. Cold Spring Harbor perspectives in biology 4

14. Wortel, I. M. N., van der Meer, L. T., Kilberg, M. S., and van Leeuwen, F. N. (2017) Surviving Stress: Modulation of ATF4-Mediated Stress Responses in Normal and Malignant Cells. Trends in endocrinology and metabolism: TEM 28, 794–806

15. Rutkowski, D. T., and Kaufman, R. J. (2003) All roads lead to ATF4. Developmental cell 4, 442–444

16. Kochetov, A. V., Ahmad, S., Ivanisenko, V., Volkova, O. A., Kolchanov, N. A., and Sarai, A. (2008) uORFs, reinitiation and alternative translation start sites in human mRNAs. FEBS letters 582, 1293–1297

17. Vattem, K. M., and Wek, R. C. (2004) Reinitiation involving upstream ORFs regulates ATF4 mRNA translation in mammalian cells. Proceedings of the National Academy of Sciences of the United States of America 101, 11269–11274

18. Wek, R. C., Jiang, H. Y., and Anthony, T. G. (2006) Coping with stress: eIF2 kinases and translational control. Biochemical Society transactions 34, 7–11

19. Saito, A., Ochiai, K., Kondo, S., Tsumagari, K., Murakami, T., Cavener, D. R., and Imaizumi, K. (2011) Endoplasmic reticulum stress response mediated by the PERK-eIF2(alpha)-ATF4 pathway is involved in osteoblast differentiation induced by BMP2. The Journal of biological chemistry 286, 4809–4818

20. Harding, H. P., Novoa, I., Zhang, Y., Zeng, H., Wek, R., Schapira, M., and Ron, D. (2000) Regulated translation initiation controls stress-induced gene expression in mammalian cells. Molecular cell 6, 1099–1108

21. Zhou, J., Wan, J., Shu, X. E., Mao, Y., Liu, X. M., Yuan, X., Zhang, X., Hess, M. E., Bruning, J. C., and Qian, S. B. (2018) N(6)-Methyladenosine Guides mRNA Alternative Translation during Integrated Stress Response. Molecular cell 69, 636–647 e637

22. Martin, J. L., and McMillan, F. M. (2002) SAM (dependent) I AM: the S-adenosylmethionine-dependent methyltransferase fold. Current opinion in structural biology 12, 783–793

23. Schubert, H. L., Blumenthal, R. M., and Cheng, X. (2003) Many paths to methyltransfer: a chronicle of convergence. Trends in biochemical sciences 28, 329–335

24. Iyer, L. M., Zhang, D., and Aravind, L. (2016) Adenine methylation in eukaryotes: Apprehending the complex evolutionary history and functional potential of an epigenetic modification. BioEssays: news and reviews in molecular, cellular and developmental biology 38, 27–40

25. Malone, T., Blumenthal, R. M., and Cheng, X. (1995) Structure-guided analysis reveals nine sequence motifs conserved among DNA amino-methyltransferases, and suggests a catalytic mechanism for these enzymes. Journal of molecular biology 253, 618–632

26. Wang, X., Feng, J., Xue, Y., Guan, Z., Zhang, D., Liu, Z., Gong, Z., Wang, Q., Huang, J., Tang, C., Zou, T., and Yin, P. (2016) Structural basis of N(6)-adenosine methylation by the METTL3-METTL14 complex. Nature 534, 575–578

27. Wang, P., Doxtader, K. A., and Nam, Y. (2016) Structural Basis for Cooperative Function of Mettl3 and Mettl14 Methyltransferases. Molecular cell 63, 306–317

28. Zorbas, C., Nicolas, E., Wacheul, L., Huvelle, E., Heurgue-Hamard, V., and Lafontaine, D. L. (2015) The human 18S rRNA base methyltransferases DIMT1L and WBSCR22-TRMT112 but not rRNA modification are required for ribosome biogenesis. Molecular biology of the cell 26, 2080–2095

29. Figaro, S., Scrima, N., Buckingham, R. H., and Heurgue-Hamard, V. (2008) HemK2 protein, encoded on human chromosome 21, methylates translation termination factor eRF1. FEBS letters 582, 2352–2356

30. Fu, D., Brophy, J. A., Chan, C. T., Atmore, K. A., Begley, U., Paules, R. S., Dedon, P. C., Begley, T. J., and Samson, L. D. (2010) Human AlkB homolog ABH8 Is a tRNA methyltransferase required for wobble uridine modification and DNA damage survival. Molecular and cellular biology 30, 2449–2459

31. van Tran, N., Ernst, F. G. M., Hawley, B. R., Zorbas, C., Ulryck, N., Hackert, P., Bohnsack, K. E., Bohnsack, M. T., Jaffrey, S. R., Graille, M., and Lafontaine, D. L. J. (2019) The human 18S rRNA m6A methyltransferase METTL5 is stabilized by TRMT112. Nucleic acids research

32. Van Nostrand, E. L., Pratt, G. A., Shishkin, A. A., Gelboin-Burkhart, C., Fang, M. Y., Sundararaman, B., Blue, S. M., Nguyen, T. B., Surka, C., Elkins, K., Stanton, R., Rigo, F., Guttman, M., and Yeo, G. W. (2016) Robust transcriptome-wide discovery of RNA-binding protein binding sites with enhanced CLIP (eCLIP). Nature methods 13, 508–514

33. Brar, G. A., and Weissman, J. S. (2015) Ribosome profiling reveals the what, when, where and how of protein synthesis. Nature reviews. Molecular cell biology 16, 651–664

34. Ingolia, N. T. (2014) Ribosome profiling: new views of translation, from single codons to genome scale. Nature reviews. Genetics 15, 205–213

35. Kimura, S., and Suzuki, T. (2010) Fine-tuning of the ribosomal decoding center by conserved methyl-modifications in the Escherichia coli 16S rRNA. Nucleic acids research 38, 1341–1352

36. Chen, M., Urs, M. J., Sanchez-Gonzalez, I., Olayioye, M. A., Herde, M., and Witte, C. P. (2018) m(6)A RNA Degradation Products Are Catabolized by an Evolutionarily Conserved N(6)-Methyl-AMP Deaminase in Plant and Mammalian Cells. The Plant cell 30, 1511–1522

37. Wu, B., Zhang, D., Nie, H., Shen, S., Li, Y., and Li, S. (2019) Structure of Arabidopsis thaliana N(6)-methyl-AMP deaminase ADAL with bound GMP and IMP and implications for N(6)-methyl-AMP recognition and processing. RNA biology, 1–9

38. Khatter, H., Myasnikov, A. G., Natchiar, S. K., and Klaholz, B. P. (2015) Structure of the human 80S ribosome. Nature 520, 640–645

39. Genuth, N. R., and Barna, M. (2018) The Discovery of Ribosome Heterogeneity and Its Implications for Gene Regulation and Organismal Life. Molecular cell 71, 364–374

40. Ferretti, M. B., and Karbstein, K. (2019) Does functional specialization of ribosomes really exist? Rna 25, 521–538

41. Genuth, N. R., and Barna, M. (2018) Heterogeneity and specialized functions of translation machinery: from genes to organisms. Nature reviews. Genetics 19, 431–452

42. Xue, S., and Barna, M. (2012) Specialized ribosomes: a new frontier in gene regulation and organismal biology. Nature reviews. Molecular cell biology 13, 355–369

43. Kondrashov, N., Pusic, A., Stumpf, C. R., Shimizu, K., Hsieh, A. C., Ishijima, J., Shiroishi, T., and Barna, M. (2011) Ribosome-mediated specificity in Hox mRNA translation and vertebrate tissue patterning. Cell 145, 383–397

44. Erales, J., Marchand, V., Panthu, B., Gillot, S., Belin, S., Ghayad, S. E., Garcia, M., Laforets, F., Marcel, V., Baudin-Baillieu, A., Bertin, P., Coute, Y., Adrait, A., Meyer, M., Therizols, G., Yusupov, M., Namy, O., Ohlmann, T., Motorin, Y., Catez, F., and Diaz, J. J. (2017) Evidence for rRNA 2’-O-methylation plasticity: Control of intrinsic translational capabilities of human ribosomes. Proceedings of the National Academy of Sciences of the United States of America 114, 12934–12939

45. Dey, S., Sayers, C. M., Verginadis, II, Lehman, S. L., Cheng, Y., Cerniglia, G. J., Tuttle, S. W., Feldman, M. D., Zhang, P. J., Fuchs, S. Y., Diehl, J. A., and Koumenis, C. (2015) ATF4-dependent induction of heme oxygenase 1 prevents anoikis and promotes metastasis. The Journal of clinical investigation 125, 2592–2608

46. Ye, J., Kumanova, M., Hart, L. S., Sloane, K., Zhang, H., De Panis, D. N., Bobrovnikova-Marjon, E., Diehl, J. A., Ron, D., and Koumenis, C. (2010) The GCN2-ATF4 pathway is critical for tumour cell survival and proliferation in response to nutrient deprivation. The EMBO journal 29, 2082–2096

47. Hsiao, K., Zegzouti, H., and Goueli, S. A. (2016) Methyltransferase-Glo: a universal, bioluminescent and homogenous assay for monitoring all classes of methyltransferases. Epigenomics 8, 321–339

48. Kaiser, R. W. J., Ignarski, M., Van Nostrand, E. L., Frese, C. K., Jain, M., Cukoski, S., Heinen, H., Schaechter, M., Seufert, L., Bunte, K., Frommolt, P., Keller, P., Helm, M., Bohl, K., Hohne, M., Schermer, B., Benzing, T., Hopker, K., Dieterich, C., Yeo, G. W., Muller, R. U., and Fabretti, F. (2019) A protein-RNA interaction atlas of the ribosome biogenesis factor AATF. Scientific reports 9, 11071

49. Dobin, A., Davis, C. A., Schlesinger, F., Drenkow, J., Zaleski, C., Jha, S., Batut, P., Chaisson, M., and Gingeras, T. R. (2013) STAR: ultrafast universal RNA-seq aligner. Bioinformatics 29, 15–21

50. Liao, Y., Smyth, G. K., and Shi, W. (2014) featureCounts: an efficient general purpose program for assigning sequence reads to genomic features. Bioinformatics 30, 923–930

51. Liu, Q., Ding, C., Chu, Y., Zhang, W., Guo, G., Chen, J., and Su, X. (2017) Pln24NT: a web resource for plant 24-nt siRNA producing loci. Bioinformatics 33, 2065–2067

52. Chan, P. P., and Lowe, T. M. (2016) GtRNAdb 2.0: an expanded database of transfer RNA genes identified in complete and draft genomes. Nucleic acids research 44, D184–189

53. Zinshteyn, B., and Gilbert, W. V. (2013) Loss of a conserved tRNA anticodon modification perturbs cellular signaling. PLoS genetics 9, e1003675

54. Langmead, B., Trapnell, C., Pop, M., and Salzberg, S. L. (2009) Ultrafast and memory-efficient alignment of short DNA sequences to the human genome. Genome biology 10, R25

55. Huang da, W., Sherman, B. T., and Lempicki, R. A. (2009) Systematic and integrative analysis of large gene lists using DAVID bioinformatics resources. Nature protocols 4, 44–57

56. Subramanian, A., Tamayo, P., Mootha, V. K., Mukherjee, S., Ebert, B. L., Gillette, M. A., Paulovich, A., Pomeroy, S. L., Golub, T. R., Lander, E. S., and Mesirov, J. P. (2005) Gene set enrichment analysis: a knowledge-based approach for interpreting genome-wide expression profiles. Proceedings of the National Academy of Sciences of the United States of America 102, 15545–15550

57. Ignatova, V. V., Stolz, P., Kaiser, S., Gustafsson, T. H., Lastres, P. R., Sanz-Moreno, A., Cho, Y. L., Amarie, O. V., Aguilar-Pimentel, A., Klein-Rodewald, T., Calzada-Wack, J., Becker, L., Marschall, S., Kraiger, M., Garrett, L., Seisenberger, C., Holter, S. M., Borland, K., Van De Logt, E., Jansen, P., Baltissen, M. P., Valenta, M., Vermeulen, M., Wurst, W., Gailus-Durner, V., Fuchs, H., de Angelis, M. H., Rando, O. J., Kellner, S. M., Bultmann, S., and Schneider, R. (2020) The rRNA m(6)A methyltransferase METTL5 is involved in pluripotency and developmental programs. Genes & development 2020 Mar 26. doi: 10.1101/gad.333369.119

58. Liberman, N., O’Brown, Z. K., Earl, A. S., Boulias, K., Gerashchenko, M. V., Wang, S. Y., Fritsche, C., Fady, P.-E., Dong, A., Gladyshev, V. N., and Greer, E. L. (2020) N6-adenosine methylation of ribosomal RNA affects lipid oxidation and stress resistance. Science Advances 2020 Apr 22: Vol. 6, no. 17, eaaz4370. doi: 10.1126/sciadv.aaz4370

